# Structures of the essential *Mycoplasma pneumoniae* lipoproteins Mpn444 and Mpn436 reveal a peptidyl-prolyl isomerase domain involved in extracellular protein folding

**DOI:** 10.1101/2024.09.05.611430

**Authors:** Irem Keles, Sina Manger, Pauline Roth, Margot P. Scheffer, Achilleas S. Frangakis

## Abstract

*Mycoplasma pneumoniae* is a human pathogen causing atypical community-acquired pneumonia. It serves as a model for a minimal cell and is notable for its absent cell wall, minimal genome, and use of antigenic variation to evade the host immune response. Here, we report the structures of the essential surface proteins Mpn444 and Mpn436 at 3.74 Å and 3.65 Å resolution, and the molecular architecture of the Mpn444 homotrimeric complex. We show that both proteins include a peptidyl-prolyl isomerase (PPIase) domain and a chaperone-like domain. *In vitro* PPIase activity assays suggest that Mpn444 and Mpn436 function as extracellular foldases in *Mycoplasma* species. Furthermore, both proteins are conserved across multiple *Mycoplasma* species. We built a composite model integrating previously reported interactions from crosslinking and cryo-ET data and we conclude that Mpn444 is responsible for the extracellular folding of nascent protein chains. Our work underscores the potential of Mpn444 and Mpn436 as a target for the development of novel strategies to treat mycoplasma infections.

**Author Summary:** In this study, we discovered that the essential proteins Mpn444 and Mpn436 of *Mycoplasma pneumoniae* act as extracellular PPIases. We solved the structures of Mpn444, Mpn436, and Mpn444’s homotrimer by single-particle cryo-EM analysis, marking them as the first experimentally resolved lipoproteins from *Mycoplasma pneumoniae*. The structures contain a peptidyl-prolyl isomerase (PPIase) domain as well as a chaperone-like domain, and the proteins show PPIase activity and bind unfolded proteins *in vitro*. By combining our structural data with previously published cryo-electron tomography and crosslinking data from *M. pneumoniae* cells, we suggest that Mpn444, and potentially Mpn436, works in concert with the Sec translocon and the membrane-bound ribosome. This finding is particularly important as the extracellular folding of bacterial proteins has been a significant gap in our understanding.

## Introduction

*Mycoplasma pneumoniae* is a human pathogen responsible for up to 40% of global cases of atypical community-acquired pneumonia. While mycoplasma infections can be treated by macrolide antibiotics, macrolide resistance is on the rise and there are no approved mycoplasma vaccines[1]. *Mycoplasmas* are minimal-cell model organisms characterized by the absence of a cell wall and the smallest known genome of any self-replicating organism[2]. *M. pneumoniae* encodes 688 ORFs and lacks the genes needed for essential metabolic pathways such as biosynthesis of the cell wall, amino acids and nucleotides[3, 4]. Furthermore, many *Mycoplasma* proteins are multifunctional: for example, some cytosolic proteins moonlight as cell adhesion proteins during host invasion[5–7]. Some cell surface proteins also undergo proteolytic processing into discrete functional domains, a process known as ectodomain shedding, by which auxiliary or secondary adhesins for host cell factors are created. Such cleavage events have been mapped in detail for Mpn142, a component of the major adhesion complex of *M. pneumoniae*, and reported for several uncharacterized lipoproteins[4]. Surface protein antigenic variation is an immune evasion mechanism used by *M. pneumoniae*. This mechanism has been reported for the major adhesion complex and several lipoproteins in other *Mycoplasma* species[8–10].

Mpn444 and Mpn436 are essential *M. pneumoniae* lipoproteins of previously unknown function which have an approximate predicted molecular weight of 145 kDa and 135 kDa, respectively[11]. Both proteins carry a lipoprotein signal sequence at the N-terminus, directly followed by a conserved cysteine that can be lipidated (as an N-palmitoyl cysteine or a S-diacylglycerol cysteine) to anchor the proteins to the mycoplasma membrane[12]. The genes *mpn444* and *mpn436* are two of 46 lipoprotein genes in six multigene lipoprotein families of *M. pneumoniae*, all of which have unknown structure and function[10]. Among this group, only five proteins, including Mpn444 and Mpn436, are essential[11]. Mpn436 is both a paralog and the most similar homolog of Mpn444 in *M. pneumoniae*, with 24% sequence identity[10]. Strikingly, the two proteins’ structure predictions appear highly similar. Given the need for highly efficient gene use in *Mycoplasma* species[13], the presence of multiple genes encoding structurally similar proteins suggests a specialized function and warrants further investigation.

Mpn444 undergoes ectodomain shedding[4], but the process has not been mapped and requires further study. It also contains short peptide variable-number tandem repeats (VNTRs)[14], which the bacteria use to evade the host immune response. Its homolog in *M. genitalium*, the uncharacterized lipoprotein MG309, has been shown to activate the host innate immune response[15]. *mpn444* shares an operon with *deaD*, an RNA helicase associated with ribosome assembly[11]. Functional studies have shown that Mpn444 can be cross-linked to the membrane protein SecD as well as Mpn436 and another uncharacterized lipoprotein, Mpn489 [16]. SecD is a homolog of the Sec translocon-associated protein SecDF, which assists in translocating unfolded polypeptide chains to the periplasmic space in multitude prokaryotes. Mpn489 is another member of the small group of essential lipoproteins and is a paralog of Mpn444[11, 17].

Mpn436 was shown to crosslink with both Mpn444 and Mpn489, as well as with another essential but uncharacterized lipoprotein, Mpn523. Additionally, both Mpn444 and Mpn436 crosslink with Mpn376, an uncharacterized protein that exhibits extensive crosslinking with numerous membrane-associated proteins[11, 16].

Here, we present the cryo-electron microscopy (cryo-EM) structure of Mpn444 at 3.74 Å resolution, and the cryo-EM structure of Mpn436 at 3.65 Å resolution, the first experimental structures of *M. pneumoniae* lipoproteins. We also identify both proteins as unfolded protein-binding PPIases based on structural similarity and *in vitro* assays. Finally, we combine previously reported crosslinking mass spectrometry and cryo-electron tomography data to create a composite model of Mpn444 with the Sec translocon and the translating ribosome. Taken together, our findings show that Mpn444 is responsible for the extracellular folding of nascent protein chains that emerge from the Sec translocon.

## Results

### Cryo-EM structures of Mpn444 and Mpn436

To investigate the structure and function of Mpn444 and Mpn436, we analyzed recombinantly expressed versions of the proteins, produced in *E. coli* without their native N-terminal signal peptides. As the proteins displayed very strong orientation bias, we used different detergents and acquired datasets with a fixed tilted stage at 0°, 20°, 30° and 40° for Mpn444, and 0° and 30° for Mpn436, to achieve sufficient angular sampling (SCF* 0.880 for Mpn444 and SCF* 0.967 for Mpn436, Supplemental Figure 1, Supplemental Figure 2). Iterative particle classification and orientation rebalancing in 2D and 3D enabled us to reconstruct the Mpn444 monomeric structure at 3.74 Å resolution (EMD-55147, Figure 1a, Supplemental Table 1, Supplemental Figure 1i - j), and the Mpn436 monomeric structure at 3.65 Å resolution (EMD-55148, Figure 1b, Supplemental Table 1, Supplemental Figure 2f - g). The density maps, which resemble the shape of Africa, span approximately 120 Å and 90 Å along the long and short axes, respectively.

**Figure 1:**
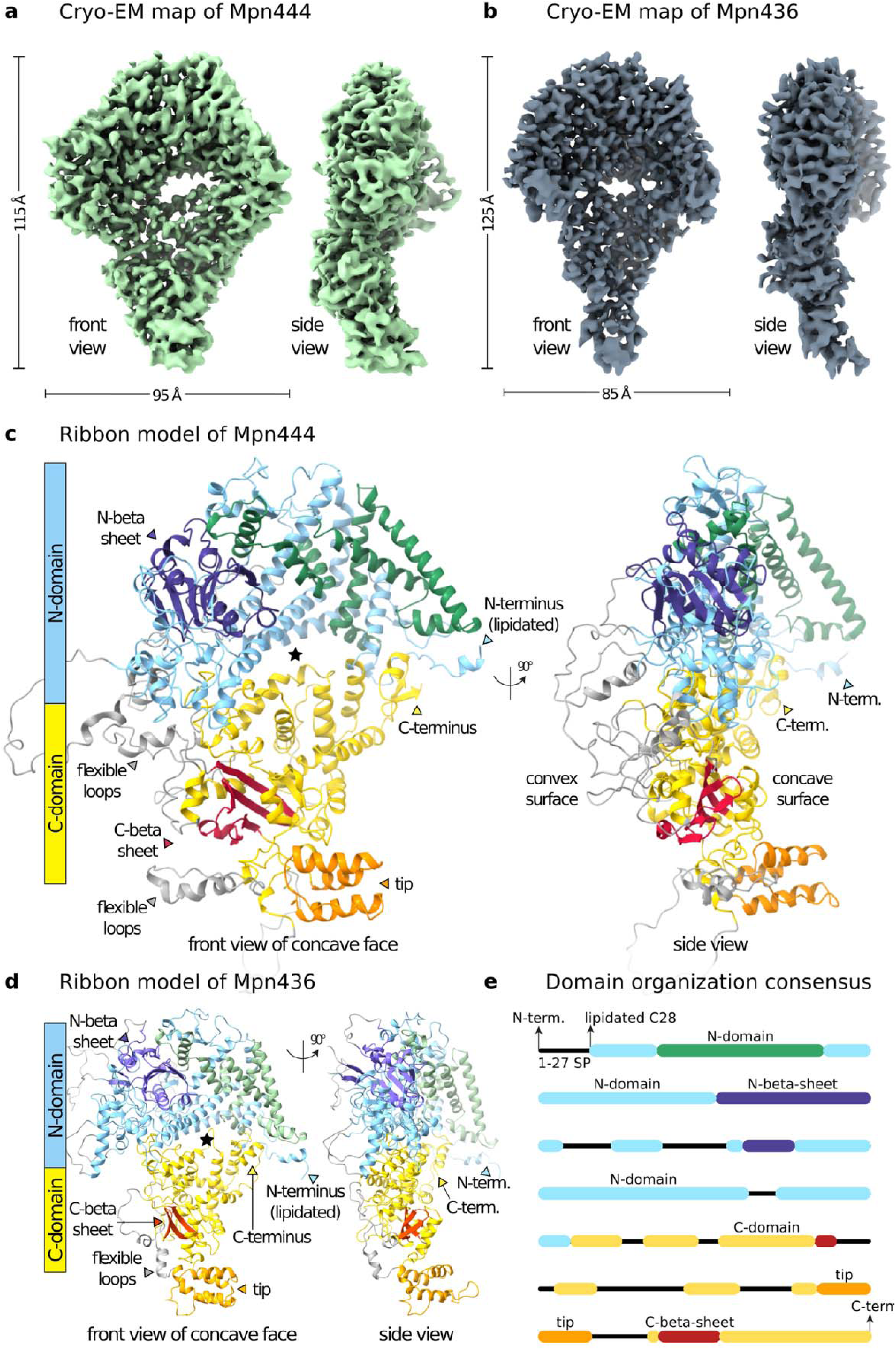
Single-particle cryo-EM of Mpn444 monomer and Mpn436. **a** Cryo-EM density map of the Mpn444 monomer at 3.74 Å. Green surface of the front view of the concave face and the side view. **b** Cryo-EM density map of the Mpn436 monomer at 3.65 Å. Slate grey surface of the front view of the concave face and the side view. **c** Front view of the concave face of Mpn444 with the structure shown as a ribbon model. The N-domain is colored blue and green (aa 54-160,514-566), the C-domain is colored yellow. The cleft separating the domains is indicated by an asterisk. The C-terminus is indicated by a yellow arrow, and the tip of the cone structure formed by the C-domain is colored orange. A small beta sheet located in the N-domain is colored purple (aa 279-352, 471-504, 230-237), and the beta sheet at the tip is colored dark red (aa 904-912,1212-1244). The lipoprotein signal peptide of Mpn444 on the N-terminus is truncated. Approximately 20% of the model consists of flexible loops with no discernable density, shown here in grey (Supplemental Figure 3). **d** Front view of the concave face of Mpn436 with the structure shown as a ribbon model. The N-domain is colored blue and light green (aa 55-158,574-626), the C-domain is colored yellow. The cleft separating the domains is indicated by an asterisk. The C-terminus is indicated by a yellow arrow, and the tip of the cone structure formed by the C-domain is colored orange. A small beta sheet located in the N-domain is colored light purple (aa 224-232,276-359,476-479,514-540), and the beta sheet at the tip is colored red (aa 956-961,1164-1178). The lipoprotein signal peptide of Mpn436 on the N-terminus is truncated. Approximately 20% of the model consists of flexible loops with no discernable density, shown here in grey (Supplemental Figure 3). **e** Schematic of the Mpn444 and Mpn436 sequence colored by domain. The N-domain is shown in blue and green, the N-beta sheet in purple, and the C-domain in yellow, the C-beta sheet in red, the tip in orange and flexible loops in black. The 27-residue signal peptide at the N-terminus was cleaved before expression.

The atomic models of Mpn444 and Mpn436 were built using the AlphaFold3 predictions as the starting models (Supplemental Table 2) and refined in Phenix and Coot to adapt them to the cryo-EM maps. We accounted for ∼80% of the sequences in the cryo-EM maps, while ∼20% of the sequence is on flexible loop regions with no clear density (Figure 1c-d, Supplemental Figure 3). The structures contain distinct N-terminal and C-terminal regions separated by a cleft (Figure 1c-d, asterisk, front view). The structures have opposing convex and concave faces with both the N- and C-termini located on the concave face (Figure 1c-d, side view). Atomic models were generated by optimizing stereochemistry through iterative manual rotamer adjustments in Coot, followed by refinement in Phenix. For each protein, we constructed two models: a full-length model (excluding only the N-terminal signal peptide) and a truncated model omitting loops not resolved in the density map (see Supplemental Table 2).

For Mpn444, the final full-length model had an atom inclusion of 0.75 and a Q-score of 0.37 (PDB: 9SRQ), with a Clashscore of 1, 1.2% sidechain outliers, and 0.1% Ramachandran outliers. The incomplete model showed improved atom inclusion (0.85) and Q-score (0.40) (PDB: 9SRS), but a higher Clashscore of 8, with 1.5% sidechain outliers and 0.5% Ramachandran outliers.

For Mpn436, the full-length model had an atom inclusion of 0.68 and a Q-score of 0.35 (PDB: 9SRR), with a Clashscore of 1, 1.2% sidechain outliers, and 0.4% Ramachandran outliers. The incomplete model exhibited higher atom inclusion (0.86) and Q-score (0.41) (PDB: 9SRV), with a Clashscore of 1, 1.7% sidechain outliers, and 0.5% Ramachandran outliers.

Sequence alignment revealed that sequences similar to the flexible loop regions of Mpn444 and Mpn436 are absent in closely related *Mycoplasma* species. In the remaining aligned regions, residues within the stable fold are notably more conserved (12% identity, 31% conservation for Mpn444, and 9% identity, 25% conservation for Mpn436) than those in the flexible loop regions (1% identity, 2% conservation for Mpn444, and 1% identity, 4% conservation for Mpn436) (Supplemental Table 3, Supplemental Figure 4, Supplemental Figure 5)[18].

A structure similarity search using the entire fold yielded no relevant results in the Protein Data Bank, although AlphaFold3[19] predicts multiple similar structures in *Mycoplasma* species, such as the essential *M. pneumoniae* lipoprotein Mpn489, its essential *M. genitalium* homolog (MG338), and the *M. genitalium* homolog of Mpn444 and Mpn436 (MG309 and MG307), as well as homologs from other *Mycoplasma* species (Supplemental Figure 6)[13, 17].

### Mpn444 and Mpn436 contain a PPIase domain and a chaperone-like domain

Since the fold search performed on the full-length Mpn444 structure did not return significant homology, we generated truncated structural models and performed independent fold searches on these models. Structural homology searches of the truncated structures revealed that the beta sheet of the N-domain, present in both Mpn444 and Mpn436, contains a peptidylprolyl isomerase (PPIase) domain (Figure 1c purple, Figure 1d light purple, Figure 2a). PPIases are enzymes that catalyze the cis-to-trans isomerization of peptide bonds N-terminal to proline residues in polypeptides, a critical step in protein folding. The domain, which contains four antiparallel β-strands and two α-helices, is representative of PPIases from the parvulin family (Figure 2a, right)[20]. Indeed, the catalytic residues in the human PPIase Pin1, the most prominent member of the parvulin family, are either conserved in Mpn444 (Pin1 S111 and H157 correspond to Mpn444 S308 and H494, respectively) or substituted with residues shown to promote full enzymatic activity in Pin1 point mutants (Pin1 C113D corresponds to Mpn444 D310)[21] (Figure 2a, left). In Mpn436, only one of the catalytic residues appears to be conserved (Pin1 S111 corresponds to Mpn436 S326). The histidine is substituted with a threonine (Pin1 H157 corresponds to Mpn436 T530). While no experimental data on Pin1 is available for this substitution, a mutation to the closely related amino acid asparagine proved functional[21]. At the position corresponding to Pin1 C113 (D310 in Mpn444), Mpn436 features a threonine; however, an aspartic acid is located just one residue away on the same loop (Figure 2a, middle). This variant is also observed in other catalytically active parvulins, such as PrsA from *Bacillus subtilis* (Figure 2a, right). The entire PPIase domains including the catalytic residues are highly conserved in Mpn444 and Mpn436 homologues from closely related *Mycoplasma* species (35% identity and 71% conservation of the PPIase domain compared to 12% identity and 31% conservation of all modelled residues for Mpn444; 15% identity and 43% conservation of the PPIase domain compared to 9% identity and 25% conservation of all modelled residues for Mpn436) (Supplemental Table 3, Supplemental Figure 4, Supplemental Figure 5).

**Figure 2:**
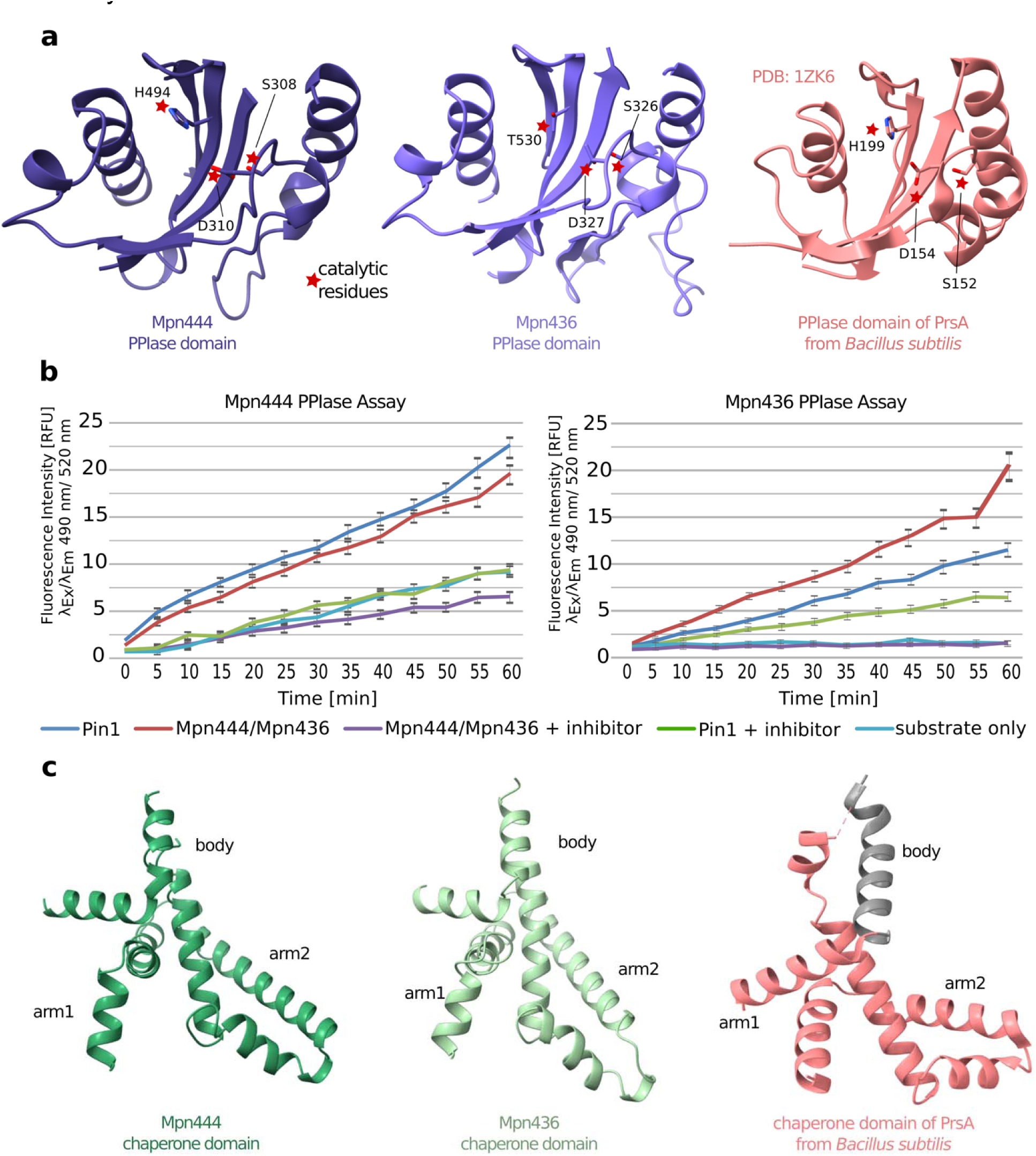
Domain functionality of Mpn444 and Mpn436. **a** The PPIase domains of Mpn444 (left, purple, aa 279-352, 471-504, 230-237), and Mpn436 (middle, light purple, aa 224-232, 276-359, 476-479, 514-540) with red stars indicating the position of the three catalytic residues of Pin1 that are conserved in Mpn444 (S282, H468 and D284), and partialy conserved in Mpn436 (T530, S328, D327), as well as the PPIase domain of PrsA from Bacillus subtilis (right, PDB: 1ZK6) with its catalytic residues (S152, H199, D154). **b** PPIase activity assay. The increase in fluorescence intensity is directly proportional to the catalyzed proline cis to trans isomerization reaction. From left to right: Pin1 (dark blue), Mpn444 or Mpn436 (red, 1 µg/µl), Pin1 with a Pin1 inhibitor (green), Mpn444 or Mpn436 with a Pin1 inhibitor (light blue) and substrate only (purple, showing the spontaneous isomerization rate of the substrate). The assay was done in three biological replicates and 35 technical replicates. **c** The chaperone-like domains of Mpn444 (green, aa 28-134, 515-540), Mpn436 (light green, aa 55-158,574-626), and of PrsA from B. subtilis (pink). The helix shown in grey is not conserved.

Parvulins are divided into a phosphor-(Ser/Thr)-Pro specific group with very high catalytic efficiency, such as human Pin1, and an unspecific group with lower catalytic efficiency, such as human Par14 or *B. subtilis* PrsA[22]. Notably, the substrate binding loop of the Pin1 PPIase domain is missing in Mpn444 and Mpn436 (Supplemental Figure 7). This loop mediates substrate recognition by specifically binding phosphorylated peptides (phosphor-(Ser/Thr)-Pro). Thus, we infer that Mpn444 and Mpn436 contain an unspecific parvulin PPIase domain. Consequently, we tested their PPIase activity with an *in vitro* assay, where cis/trans substrate peptide isomerization led to fluorescence emission. Both Mpn444 and Mpn436 demonstrated PPIase activity, which was abolished by the addition of a Pin1 inhibitor (Figure 2b).

We further investigated the PPIase activity of Mpn444 by introducing a single-point mutation at the catalytic site (D310). Substitution of aspartate with alanine (Mpn444-D310A) resulted in a reduction in PPIase activity, while the overall protein fold remained intact as determined by cryo-EM analysis (Supplemental Figure 8). In contrast, the corresponding mutant of Mpn436 (D327A) did not express under the tested conditions, despite multiple attempts, which may reflect differences in stability or folding requirements between the two proteins. Additional targeted mutants are currently being generated to further evaluate the contribution of other residues within the catalytic region to PPIase activity.

In addition to the parvulin PPIase domain, we also identified a chaperone-like domain (Figure 1c green, Figure 1d light green, Figure 2c). Despite being unrelated in sequence, its fold is structurally related to the chaperone domains of the trigger factor and SurA from *E. coli*, PrsA from *B. subtilis* and LIC12922 from *Leptospira interrogans*, which are organized in a characteristic ‘arms’ and ‘body’ structure (Figure 2c). In these proteins, the body typically serves as a scaffold for substrate binding, while the arms assist in substrate capture and stabilization[23, 24]. In all four proteins, a PPIase domain is located directly adjacent to the chaperone domain[24], which is a common feature in foldases, where the two domains work together to facilitate protein folding and stabilization[25]. To assess the chaperone activity of Mpn444 and Mpn436, we performed two complementary assays: (1) evaluating their ability to prevent unfolding and aggregation of a client protein, and (2) testing their capacity to selectively bind unfolded clients.

For the first assay, alcohol dehydrogenase (ADH) was heat-inactivated, and recovery of its catalytic activity was monitored in the presence and absence of Mpn444 and Mpn436[26]. Under these conditions, ADH activity could not be restored, indicating that Mpn444 and Mpn436 were unable to refold the denatured protein. However, this lack of activity may reflect the absence of co-factors or partner proteins that are required for chaperone function, or substrate specificity limited to *Mycoplasma*-native proteins rather than ADH.

In the second assay, heat-inactivated ADH was incubated with Mpn444 or Mpn436, and the mixtures were analyzed by size-exclusion chromatography (SEC). Both Mpn444 and Mpn436 co-eluted with heat-inactivated ADH, suggesting the formation of a complex. In contrast, no complex formation was observed between either protein and native (non-denatured) ADH. Additionally, lysozyme did not co-elute with heat-inactivated ADH, confirming the specificity of the interaction (Supplemental Figure 9). These results suggest that Mpn444 and Mpn436 function as unfolded protein-binding PPIases, possibly implying a role in early folding or interaction with other folding machinery.

### Mpn444 forms homotrimeric complexes

In the cryo-EM classification of Mpn444 particles, we routinely observed trimeric complexes (approximately 20% of particles; Supplemental Figure 10a) and dimeric complexes (approximately 20% of particles; Supplemental Figure 10b). After reconstruction without symmetry imposition, a C3 symmetric density prevailed; it had a weaker density in one monomer, likely originating from the persistent presence of monomers and dimers in the particle set even after several rounds of 2D classification (Supplemental Figure 10c). The dimers appeared to be incomplete trimeric assemblies, as they displayed the same subunit architecture as trimers. Therefore, we reconstructed the final density map using C3 symmetry and reached a global resolution of 4.5 Å (5.8 Å according to the EMDB validation) with the best areas resolved at ∼3.5 Å (PDB: 9SSC, EMD-53144; Figure 3a, Supplemental Figure 10d & e, Supplemental Table 1).

**Figure 3:**
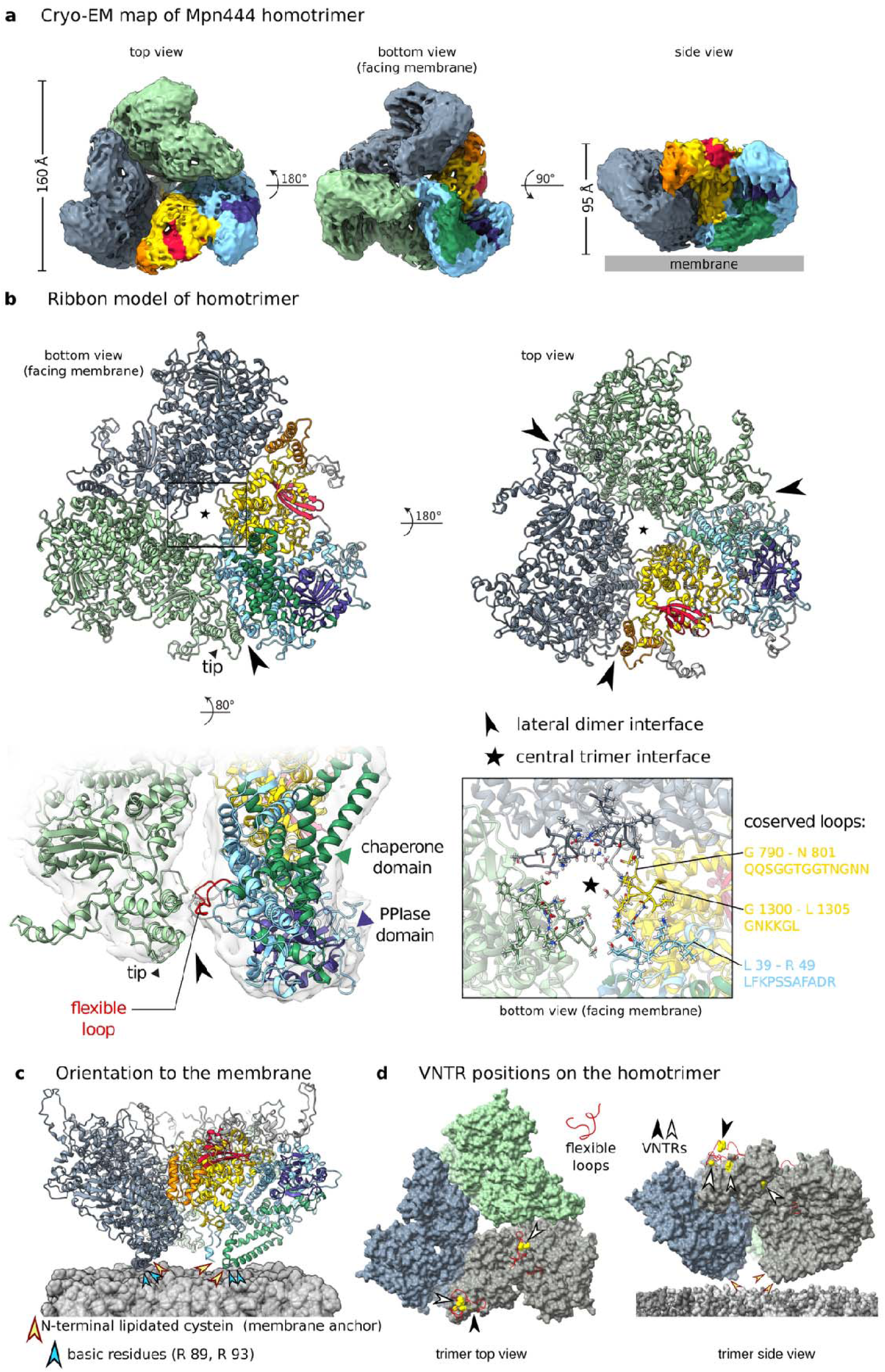
Multimeric organization of Mpn444. **a** Surface representation of the cryo-EM map of the Mpn444 homotrimer at 5.4 Å resolution as viewed from the bottom (facing the membrane), top and side with the position of the membrane indicated. The surface of one monomer is colored according to the color of the monomer ribbon models in Figure 1c. **b** Molecular architecture of the Mpn444 trimer from a rigid body fit of monomer models of Mpn444 into the cryo-EM map of the trimeric complex. The ribbon model of one monomer is shown in the domain color code as in Figure 1c. The two other monomers are shown in green and blue. The central trimer interface is indicated by an asterisk, and the lateral dimer interface is indicated by a black arrow. The lateral dimer interface (bottom left) connects the tip of one monomer to the lower convex face of the adjacent monomer, leaving the chaperone-like and PPIase domain on the concave face accessible. The loop shown here in red (Gly675–Asn691: GT**STVKTTSSNTG**QTKN) is situated in an area of the map with density that is not accounted for by the monomer structures and could be involved in trimer formation or stabilization. A channel is formed at the central trimer interface (bottom right), indicated by an asterisk, with a diameter of 23 Å on the top and 11.5 Å on the bottom composed of three short loops with conserved residues. **c** Orientation of Mpn444 to the membrane. The C-terminal lipidated cysteines that mediate membrane anchoring are indicated with yellow arrows. The basic residues R 89 and R93 on the membrane facing interface are indicated with blue arrows. **d** Positions of VNTRs[14] relative to the trimer architecture (shown on one monomer only), indicated by black or black/white arrows. The residues involved are shown as yellow spheres; VNTR positions appear to be accessible and facing away from the cell membrane.

Three Mpn444 monomers assemble into a C3 symmetric homotrimeric complex, resembling a propeller. In the center, a tapering channel is formed with a diameter of 23 Å on the top and 11.5 Å on the bottom (Figure 3b, Supplemental Figure 10c). This channel is composed of three short loops which comprise the central trimer interface and are conserved in the close homolog MG309 from *M. genitalium*. The amino acid compositions are Gln790–Asn801 (QQSGGTGGTNGNN), Gly1300–Leu1305 (GNKKGL) and Leu39–Arg49 (LFKPSSAFADR), creating a hydrophilic and moderately basic interface surface (Figure 3b). Due to resolution limitations of the trimer map, loop placements at the trimer interface are approximate and do not support functional interpretations. The lateral dimer interfaces and the central trimer interface have additional densities that are unaccounted for through rigid body docking of the monomers (Supplemental Figure 3b). Specifically, the lateral dimer interface contains a loop that could not be resolved in the monomeric structure (Gly675–Asn691: GTSTVKTTSSNTGQTKN; Figure 3b, bottom left), making it likely to be involved in trimer formation or stabilization.

As viewed from the bottom (membrane-binding side), the convex face of Mpn444 is on the inside of the complex, and the tip of one monomer links to the lower convex face of the adjacent monomer thereby forming the lateral dimer interface (Figure 3b). Both the PPIase and chaperone-like domains are located on the concave face, suggesting that trimer formation does not interfere with their catalytic activity. The N-termini of the monomers, which contain the membrane-anchoring lipidated cysteine residues, are all arranged on the bottom of the complex, unambiguously confirming its orientation towards the membrane (Figure 3c). Directly adjacent to the lipidated cysteine residues, two positively charged amino acids (Arg115 and Arg119) are positioned on the membrane-facing interface. These residues may facilitate interactions with the membrane to stabilize the complex’s orientation or potentially contribute to its membrane association. Interestingly, this architecture is completely different from the AlphaFold3 prediction of the trimer (Supplemental Figure 11).

We evaluated our homotrimeric assembly of Mpn444 in the context of available crosslinking data[16] and found that six of the seven amino acids involved in crosslinks with other *Mycoplasma* proteins are located at the periphery of the complex, making them accessible for protein-protein interactions (Supplemental Figure 12). The variable-number tandem repeats (VNTRs), through which Mpn444 contributes to host immune evasion, are also exposed at the complex periphery and face away from the membrane (Figure 3d). This arrangement likely facilitates interactions with components of the host immune system.

### Composite model of Mpn444 based on crosslinking mass-spectrometry data and cryo-electron tomography

We analyzed previously reported crosslinking mass spectrometry data on the interactions between Mpn444 and SecD[16]. SecD is a key component of the Sec translocon, responsible for translocating newly synthesized proteins into the periplasmic space in gram-negative bacteria, and to the extracellular space in the genus *Mycoplasma*. In *M. pneumoniae*[27], SecD has relatively few crosslinks overall, with Mpn444 being the only protein showing multiple interactions with three crosslinks (Supplemental Figure 12a). By contrast, SecD forms single crosslinks with other membrane proteins such as P30, which is involved in host cell adhesion, and Mpn052, an uncharacterized lipoprotein that likely functions as a BMP family ABC transporter substrate-binding protein[28]. Given its role in protein translocation, the Sec translocon is expected to interact with chaperones or peptidylprolyl isomerases in the periplasmic or extracellular space to facilitate the proper folding of newly translocated proteins. Based on our results, which show that Mpn444 is a PPIase and specifically binds unfolded proteins, the interaction between Mpn444 and SecD appears biologically relevant.

By integrating these findings, we constructed a composite model of Mpn444 in association with SecD and SecYEG. Due to the absence of experimental structures, we relied on AlphaFold predictions of the Sec-associated proteins. The predictions show a high degree of similarity to the experimental structures of homologs from other gram-negative bacteria (Supplemental Figure 13). Based on crosslinking data, we positioned SecD beneath the propeller of Mpn444 (Figure 4). The position of SecYEG, which forms the translocation channel across the membrane, and SecA, which drives translocation from the cytosolic side through ATP consumption, was determined from AlphaFold’s prediction of the complex. This complex architecture aligns with experimental structures from other Gram-negative bacteria[29–31]. The resulting model suggests that Mpn444 collaborates with SecD to facilitate the folding of nascent protein chains emerging from SecYEG (Figure 4).

**Figure 4:**
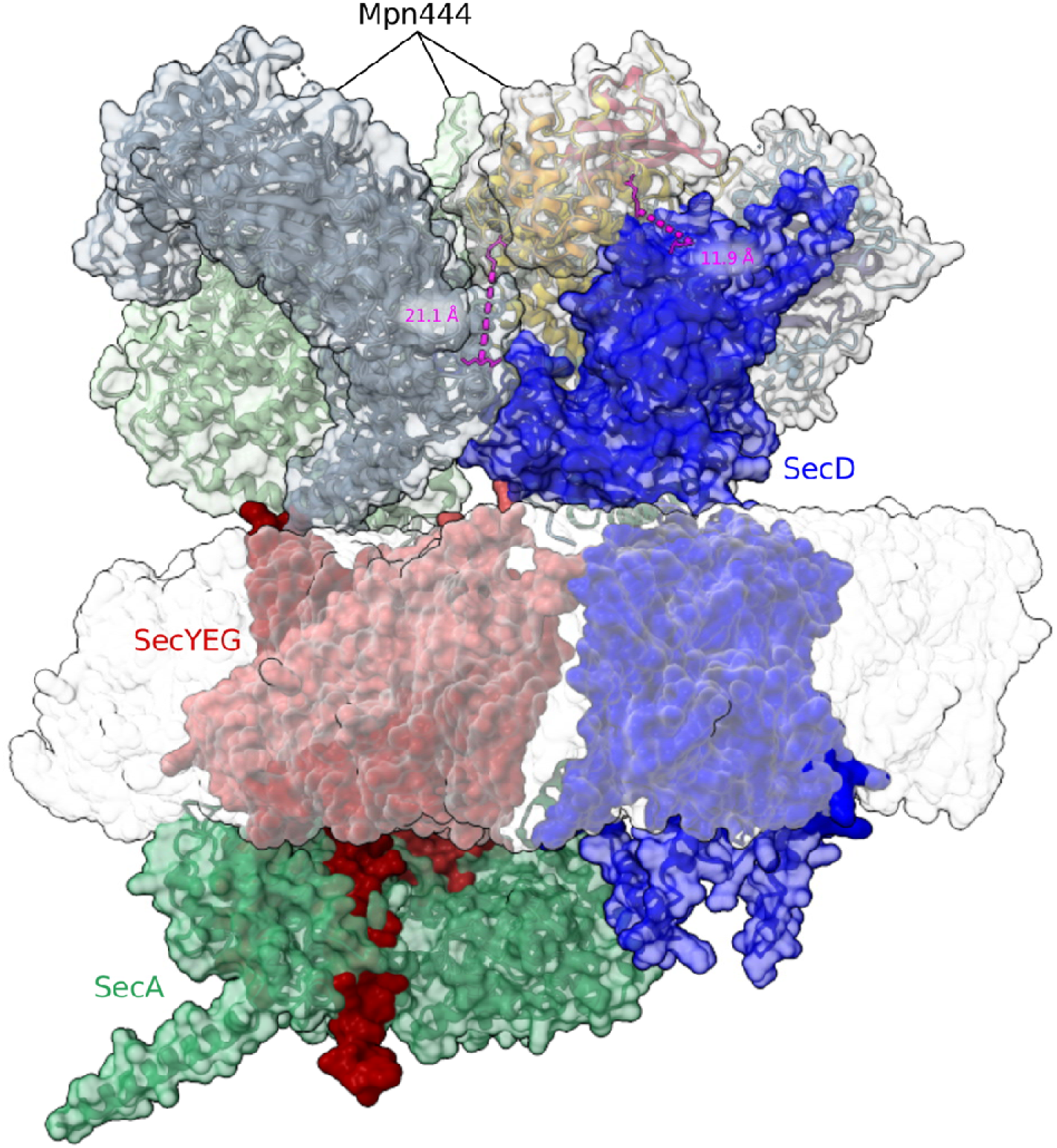
Composite model of the proposed complex of the Sec translocon and the Mpn444 trimer. SecYEG (three separate proteins in *M. pneumoniae*) is shown in red (AF-Q59548-F1-v4, AF-P75048-F1-v4, and AF-Q9EXD0-F1-v4, respectively), SecA in green (AF-P75559-F1-v), and SecD (equivalent of SecDF in *E. coli*) in blue (AF-P75387-F1-v4). The Mpn444 trimer is shown with the same color coding as previously. The crosslinks between SecD and Mpn444 are indicated in magenta. The membrane is shown as a white, transparent surface. The AlphaFold3 predictions of SecD, SecYEG, and SecA are highly similar to the structures of homologs from other gram-negative bacteria (Supplemental Figure 13).

The Sec translocon has been well-documented to form complexes with translating ribosomes[32, 33], allowing nascent proteins to be directly translocated through the SecYEG pore into the periplasmic or extracellular space, where they are folded. This implies that complexes involving ribosomes, SecYEG, and Mpn444 should exist in *Mycoplasma*.

To investigate whether Mpn444 may be involved in folding nascent protein chains emerging from the Sec translocon, we performed subtomogram averaging of ribosomes using a previously published cryo-electron tomography dataset from *M. pneumoniae* (Cryo-ET Data Portal, dataset ID: DS-10003)[34, 35]. The resulting average revealed that (i) both the ribosome and the membrane were well resolved, indicating a conserved ribosome orientation relative to the membrane consistent with a stable interaction, and (ii) a trimeric density was observed adjacent to the ribosome on the extracellular side of the membrane (Supplemental Figure 14). While the resolution of this map does not permit definitive molecular identification, the size, shape, and position of this trimeric density are consistent with our structural model of Mpn444.

This assignment is further supported by previously published crosslinking mass spectrometry data, which identified multiple crosslinks between Mpn444 and SecD—an interaction not observed with other membrane-associated proteins to the same extent. Given Mpn444’s dual chaperone-like and PPIase domains and its conservation across species, its proximity to the translocon complex is functionally plausible.

However, we emphasize that this interpretation is tentative and based on indirect evidence. Although the structural model is consistent with known ribosome–Sec–chaperone assemblies in other bacteria, definitive assignment of Mpn444 to this trimeric density will require additional experimental validation, such as *in situ* labeling or higher-resolution tomography. We therefore present this as a working model that integrates current structural and biochemical data, offering a hypothesis for future investigation.

## Discussion

Here, we present the first experimental structures of Mpn444 and Mpn436, which are extracellular PPIases and putative chaperones from *Mycoplasma pneumoniae*. We conclude that Mpn444 may act in concert with the ribosome and the Sec translocon to assist in the extracellular folding of nascent protein chains emerging from SecYEG (Figure 4). This hypothesis is based on (i) the structure of the homotrimer, (ii) its extracellular localization, (iii) biochemical assays and structure-function relationship, (iv) the previously reported interactions between Mpn444 and SecD, and (v) a sub-tomogram average showing co-localization of a trimeric density extracellularly associated with the ribosome. Importantly, our conclusion is now independently supported by a recent cryo-ET preprint that reports an in situ sub-tomogram average of a trimeric assembly associated with the Sec translocon[36]. In this study, both the Sec translocon components and the associated ribosome are clearly resolved, providing orthogonal structural evidence that supports our proposed model of Mpn444 as a folding factor linked to the bacterial secretion machinery.

Given that *Mycoplasma* species lack a periplasm, membrane proteins and other secreted proteins must fold in the extracellular space. In contrast to gram-negative bacteria, where multiple periplasmic chaperones such as SurA, Skp, PpiD and YfgM have been characterized[37–39], no extracellular foldases have been identified or characterized in *Mycoplasma* prior to this study. Comparison of our structures with AlphaFold predictions of homologous proteins suggests that the here presented fold is conserved across multiple *Mycoplasma* species (Supplemental Figure 6).

Given the structural similarity between Mpn444, Mpn436, and the essential lipoprotein Mpn489, their comparable expression levels (48, 57, and 48 copies on average per cell, respectively)[11], and the multiple crosslinks observed among them these three proteins[16], we propose that these proteins may form a heterotrimeric complex *in vivo*. This model could account for the lack of chaperone activity observed for the Mpn444 monomer and homotrimer *in vitro*. A heterotrimeric assembly would likely adopt the same overall architecture and dimensions as the observed homotrimer and would therefore be consistent with the trimeric density identified in the sub-tomogram average. Notably, the trimeric arrangement reported *in situ* in an independent cryo-ET study - although assigned by the authors to an Mpn444–Mpn436–Mpn489 heterotrimer based on indirect evidence - would likewise be compatible with either a heterotrimeric or homotrimeric complex formed by any of these individual proteins[36].

Despite their notoriety as the simplest known bacteria, there is little known regarding the translation, translocation and extracellular folding of proteins coordinated in *Mycoplasma* species. By contrast, complexes associated with the translating ribosome and the Sec translocon have been reported for a multitude of pro-and eukaryotic species[40–42]. Understanding how the ribosome, the Sec translocon, and extracellular foldases interact could provide key insights into how fundamental processes such as protein synthesis and translocation occur in the absence of a cell wall and additional complex protein transport systems. While AlphaFold predicted the monomer structure, the prediction of the homotrimer is different to our experimental structure for the trimeric complex. Further investigation into the interaction of Mpn444, and potentially Mpn436 and Mpn489, with the Sec translocon in its cellular context will be crucial for advancing our understanding of its function *in vivo*.

Finally, although *M. pneumoniae* infections are treatable with macrolide antibiotics, macrolide resistance is on the rise and no approved vaccines exist[43]. Considering that Mpn444 is an essential protein that is also involved in host immune evasion through antigenic variation, our structure presents opportunities to elucidate the pathogenesis of *M. pneumoniae* community-acquired pneumonia and to develop new treatment strategies against this pathogen.

## Material & Methods

### Expression and purification of Mpn444, Mpn444-D310A, and Mpn436

For expression, we used a pET-21(+) vector containing a codon optimized *mpn444* or *mpn436* constructs without the N-terminal signal sequence (leaving aa 28–1325 for Mpn444, aa 28-1217 for Mpn436) but with a C-terminal His6-tag (Twist Bioscience, San Francisco, CA, USA). Same protein expression conditions and purification procedures were applied for all three proteins. *E. coli* BL21 (DE3) cells harboring the vector were grown overnight at 37°C in LB medium. The culture was inoculated in 2x YT autoinduction medium supplemented with 0.2% lactose, 0.05% glucose and 0.6% glycerol and grown overnight at 37°C with mild shaking. Cells were harvested, lysed by sonication in lysis buffer (50 mM TRIS-HCl at pH 7.4, 200 mM NaCl, 20 mM imidazole, 1 mM PMSF, 0–0.5% LMNG), and centrifuged at 15,000 x g (Centrifuge 5804 R, Eppendorf, Hamburg, Germany) for 15 min at 4°C and again at 20,400 x g in a tabletop centrifuge for 30 min at 4°C. The supernatant was loaded onto a HisTrap 5 mL column (GE Healthcare, Chicago, IL, USA) that was pre-equilibrated with binding buffer (20 mM TRIS-HCl, pH 7.4, 200 mM NaCl, 20 mM imidazole, 0–0.5% LMNG), thoroughly washed with binding buffer and eluted with elution buffer (20 mM TRIS-HCl pH 7.4, 200 mM NaCl, 400 mM imidazole, 0–0.5% LMNG). The eluate was loaded onto a Superose 6 Increase 10/300 GL column (Cytivia, Marlborough, MA, USA) in protein buffer (20 mM TRIS-HCl, pH 7.4, 200 mM NaCl, 0–0.005% LMNG). The expression yield was approximately 2.5 mg/L for Mpn444, 2 mg/L for Mpn444-D310A, and 2.8 mg/L for Mpn436. Fractions were pooled and stored at –80°C. The identity and purity of Mpn444 and Mpn436 were confirmed by SDS-PAGE and mass spectrometry (carried out by the Functional Genomics Center at ETH Zurich, Switzerland).

### Cryo-EM sample preparation and data acquisition

A 3.5 μL drop of purified protein (0.2 mg/mL of Mpn444, 0.5 mg/mL of Mpn444-D310A or 0.5 – 0.8 mg/mL of Mpn436 in buffer containing 20 mM TRIS-HCl pH 7.4, 200 mM NaCl and 0–0.01% LMNG or 1 mM CHAPSO) was applied to a 45 s glow-discharged R1.2/1.3 C-flat grid (Electron Microscopy Science, Hatfield, PA, USA) or R1.2/1.3 300 copper mesh quantifoil grid (Quantifoil Micro Tools, Großlöbichau, Germany). Grids were plunge-frozen in liquid ethane (Vitrobot Mark IV; Thermo Fischer Scientific, Eindhoven) at 4°C, 100% relative humidity, with a nominal blot force of –3, a wait time of 10 s, and a blotting time of 6 - 14 s. Before freezing, Whatman 595 filter papers were incubated for 1 h in the Vitrobot chamber at 100% relative humidity and 4°C.

Dose-fractionated movies of Mpn444 were collected with SerialEM v4.1.0 beta[44] at a nominal magnification of 105,000 (0.92 Å per pixel) in nanoprobe EFTEM mode at 300 kV with a Titan Krios electron microscope (Thermo Fischer Scientific) equipped with a GIF Quantum S.E. post-column energy filter in zero-loss peak mode and a K3 Summit detector (Gatan, Pleasanton, CA, USA). The camera was operated in dose-fractionation counting mode with a dose rate of ∼16 electrons per Å^2^ s^−1^, resulting in a total dose of 50 electrons per Å^2^. The frame time was adjusted to 1 frame per 1 electron per Å^2^. A total of 7,811 for Mpn444, 4,596 for Mpn444-D310A and 6,726 for Mpn436 micrographs were recorded at defocus values ranging from –0.8 to –3.5 μm. Micrographs were acquired at 0°, 20°, 30° and 40° stage tilts for Mpn444, and 0° and 30° stage tilts for Mpn436, to overcome the evident preferred orientation of the particles.

### Cryo-EM image processing

CryoSPARC v4.5.3[45] was used to process the cryo-EM data, unless stated otherwise. Beam-induced motion correction and contrast transfer function (CTF) estimation were performed separately for each stage tilt dataset using cryoSPARC’s own implementation. For the monomers, particles were initially selected with the blob picker using a particle diameter of 100–200 Å. Particles were then subjected to unsupervised 2D classification. The selected 2D classes were used as templates to retrieve more particles from the micrographs.

For Mpn444, a total of 6,832,694 particles were extracted from 7,811 micrographs using a box size of 480 pixels and were subjected to unsupervised 2D classification. Selected 2D classes were uniformly rebalanced (using 8 superclasses and a rebalancing factor of 1). The remaining 2,992,076 particles were used to generate an *ab initio* reconstruction with 3 classes. Subsequent non-uniform refinement was followed by local and global CTF refinement and another non-uniform refinement[46]. Orientations were then rebalanced in 3D, yielding a final set of 432,841 particles. The resulting refined map was further processed by filtering map amplitudes, in which amplitudes overrepresented in the power spectrum were proportionally attenuated relative to non-overrepresented amplitudes. The final map had a global monomeric resolution of 3.74 Å (EMD-55147, Figure 1a, Supplemental Figure 1, Supplemental Table 1).

For the homotrimers, particles were initially selected with the blob picker using a particle diameter of 150–300 Å. Particles were then subjected to unsupervised 2D classification. Particles were also selected with the template picker using the generated 2D averages as templates. After removal of duplicate particles, a total of 3,376,184 particles were extracted with a box size of 480 pixels. False-positive picks were removed by two rounds of unsupervised 2D classification. The remaining 1,126,935 particles were used to generate *ab initio* reconstructions with C1 and C3 symmetry. Since non-symmetrized reconstructions displayed the same overall domain architecture as the C3-symmetrized reconstruction, all following refinement steps were performed with C3 symmetry applied.

Following non-uniform refinement[46], the particles were further classified in 3D and individual classes were refined locally. The obtained global resolution of the trimer using 506,291 particles was 4.49 Å (5.85 Å according to EMDB validation) with a resolution range of 3.5–8 Å (EMD-53144, Supplemental Figure 10, Supplemental Table 1).

For Mpn436, a total of 6,582,748 particles were extracted from 6,726 micrographs using a box size of 480 pixels and were subjected to unsupervised 2D classification. Selected 2D classes were uniformly rebalanced (using 6 superclasses and a rebalancing factor of 1). The remaining 2,168,069 particles were used to generate an *ab initio* reconstruction with 3 classes. Subsequent non-uniform refinement was followed by local and global CTF refinement and another non-uniform refinement[46]. Orientations were then rebalanced in 3D, yielding a final set of 468,211 particles. The refined consensus map at a global resolution of 3.65 Å (EMD-55129) was subjected to local refinement of the tip region, yielding a focused map at at a global resolution of 4.07 Å (EMD-55131). The composite map isotropy was further optimized by filtering map amplitudes, in which amplitudes overrepresented in the power spectrum were proportionally attenuated relative to non-overrepresented amplitudes (EMD-55148, Figure 1b, Supplemental Figure 2, Supplemental Table 1).

CryoSPARC v4.7.1 was used to process Mpn444-D310A cryo-EM data. Initially, particles were selected with blob picker and template picker with a particle diameter range of 100-200 Å. In total, 4,992,685 particles were extracted from 4,034 micrographs using a box size of 256 pixels and were subjected to unsupervised 2D classification. After removing the duplicate particles, an *ab initio* model with 2 classes was generated using 1,276,010 particles. The reconstructions were used in heterogenous refinement followed by a non-uniform refinement. Then, we perform a 3D variability analysis to assess the conformational heterogeneity. The final round of non-uniform refinement resulted in a map at a global resolution of 3.52 Å. The resulting refined map was further processed by filtering map amplitudes, in which amplitudes overrepresented in the power spectrum were proportionally attenuated relative to non-overrepresented amplitudes.

### Model building

The quality of our maps was initially assessed using ModelAngelo[47] with and without the FASTA sequence. For the full-length models, AlphaFold3-predicted structures[48] were flexibly fitted into the density maps using ISOLDE [49]. For the partial models, AlphaFold3 predictions[48] were used as input for the *predict_and_build* tool in Phenix[50] to generate an initial model. Missing loops were modeled using the *fit_loops* tool in Phenix, but only in regions with sufficient supporting map density. Loops lacking defined density were not modeled, resulting in an overall sequence coverage of approximately 80%.

Both full-length and partial models underwent iterative refinement using a combination of global real-space refinement in Phenix, per-residue refinement via the Real-Space Refinement Zone tool in Coot v0.9.8[51] and manual rotamer adjustment in Coot as needed. Final models were deposited in the Protein Data Bank (PDB) under the accession codes listed in Supplemental Table 2.

For the Mpn444-D310A mutant, we used the native Mpn444 structure as starting model and fitted into the Mpn444-D310A density map using ISOLDE. The resulting model showed a strong overall fit to the density. We further refined the catalytic site invluding D310A substitution using Real-Space Refinement Zone tool in Coot v0.9.8.

The resolution of the Mpn444 trimer cryo-EM density map was too poor to allow for *ab initio* model building. Thus, the model of the trimeric assembly was assembled by fitting the cryo-EM density map of the monomer into the cryo-EM density map of the trimer and then rigid-body fitting the monomer model into the monomer density maps. The model is deposited in the PDB under the accession code 9SSC (Supplemental Table 2). All cryo-EM density maps and models were visualized and analyzed using UCSF ChimeraX[52].

### Structural similarity search

Protein structure databases were searched using individual domains of Mpn444 and Mpn436 using FoldSeek[53] and DALI servers[54] to identify structurally similar proteins. Predicted structures of Mpn444 and Mpn436 homologs in other *Mycoplasma* species were taken from the AlphaFold Protein Structure Database[55].

### *In vitro* chaperone activity

The putative chaperone activities of Mpn444 and Mpn436 were tested with two complementary approaches: (i) evaluating the ability to recover an unfolded client by an ADH activity assay as described in a previous study[26] and (ii) assessing the selective binding to unfolded clients by size-exclusion chromatography (SEC). For the first approach, we first measured the activity of 0.1 µM yeast ADH (A7011; Sigma, St Louis, MO, USA) at 340 nm for 3 min at room temperature immediately after the addition of the final concentration of 6 mM NAD^+^ (N7004; Sigma) and 3.75 mM ethanol. Then, we deactivated ADH at 46°C for 30 min and 60 min. To test the chaperone activity of Mpn444 and Mpn436 at concentrations of 0.1 µM, 0.5 µM, and 1 µM, two separate reactions were conducted, with Mpn444 or Mpn436 added either before or after heat inactivation. The recovery of ADH activity after heat inactivation was observed at 340 nm for 3 min.

For the second approach, we first observe the elution profiles of native ADH and denatured ADH that had been heat-inactivated at 46°C for 1 h as described above. We then incubated denatured ADH with either Mpn444 or Mpn436 at a 1:1 (w/w) ratio on ice for 20 min prior to loading onto Superose 6 Increase 10/300 GL column (Cytiva, Marlborough, MA, USA). Proteins were eluted in 20 mM TRIS-HCl, pH 7.4 with 200 mM NaCl. To assess non-specific interaction, we also conducted gel filtration of Mpn444 or Mpn436 with native ADH as well as lysozyme with denatured ADH.

### *In vitro* PPIase activity

The PPIase activities of Mpn444, Mpn444-D310A, and Mpn436 were determined independently using a SensoLyte Green Pin1 Activity Assay Fluorimetric kit (AS-72240; AnaSpec, Fremont, CA, USA). According to the manufacturer’s protocol, 2.5 µg of Mpn444 or Mpn436 were incubated with 1 µM substrate for 30 min. Fluorescence was measured for 60 minutes using a CLARIOstar Plus plate reader (BMG Labtech, Ortenberg, Germany) at an excitation wavelength of 490 nm and an emission wavelength of 520 nm. As a positive control, 500 ng of human Pin1 enzyme, provided in the kit, was used. Additionally, the effect of inhibition on Mpn444, Mpn436 and Pin1 activity was evaluated using 100 µM of the inhibitor provided in the kit.

### Sub-tomogram averaging

For sub-tomogram averaging (STA) of membrane-bound ribosomes, we used a dataset of M. pneumoniae cells annotated with ribosome positions from the CryoET Data Portal[34] (dataset ID: DS-10003)[35]. First, we classified the ribosomes by their distance to the mycoplasma membrane, using customized matlab scripts. Ribosomes within a distance of 28 px of the membrane (1992 sub-tomograms at a pixel size of 6.802 Å) were then averaged with the STA routine of Artiatomi (https://github.com/uermel/Artiatomi)[56, 57], using a box size of 96 px. A first round of STA revealed a ribosome bound to the membrane with a small extracellular density co-localizing with the ribosome. We then performed a second round of STA, using a mask to exclude the ribosome density and thereby aligning on the extracellular density. This STA revealed a trimeric density that matched the size of the Mpn444 homotrimer. Finally, we applied C3-symmetry during STA to produce the final map of the extracellular density.

## Data availability

The cryo-EM structures solved in this study are available in public databases under the accession numbers EMD-55147, EMD-53144, EMD-55148, PDB: 9SRQ, PDB: 9SRS, PDB: 9SSC, PDB: 9SRR, and PDB: 9SRV. The vectors are available upon request.

## Acknowledgements and funding

We thank the Frankfurt Center for Electron Microscopy for measurement time. We thank the Pos lab (Goethe University Frankfurt) for their support with the ÅKTA system. A. S. F. acknowledges support from the Deutsche Forschungsgemeinschaft and the Research Training Group iMOL (GRK 2566/1 for S.M. and I.K.). The funders had no role in study design, data collection and analysis, decision to publish, or preparation of the manuscript.

## Author contributions

I.K.: Expressed and purified Mpn444 and Mpn436, prepared samples for single-particle analysis, performed *in vitro* activity assays. S.M.: Processed single-particle datasets of Mpn444 monomers and trimers, and Mpn436 monomers, built and refined the molecular model of Mpn444 and Mpn436, wrote the manuscript with contributions from all authors. I.K. and S.M.: Designed the expression plasmid for Mpn444 and Mpn436, recorded single-particle datasets of Mpn444 and Mpn436. P.R.: Did the sub-tomogram averaging. M.P.S.: Designed the research. A.S.F.: Designed and supervised the research.

## Supporting Information

**Supplemental Table 1: Single-particle cryo-EM data collection and processing**

**Supplemental Table 2: Model building statistics**

**Supplemental Table 3: Conservation of modelled and unmodelled regions**

Statistical evaluation of a sequence alignment of Mpn444 or Mpn436 from *Mycoplasma pneumoniae* with homologs from five other *Mycoplasma* species (*M. genitalium*, *M. bradburyae*, *M. gallisepticum, M. testudineum*, *M. amphoriforme*) (Supplemental Figure 4, Supplemental Figure 5). Residues within the stable fold are significantly more conserved (12% identity, 31% conservation for Mpn444, and 9% identity, 25% conservation for Mpn436) than residues on flexible loops (1% identity, 2% conservation for Mpn444, and 1% identity, 4% conservation for Mpn436). ‘Conserved’ includes amino acids with a >5 conservation score. ‘Identical’ includes residues that are unchanged across all homologs (labelled with a (9)). The total residue count excludes the N-terminal signal peptide, which was not part of the structure.

**Supplemental Figure 1: Single-particle cryo-EM of Mpn444 monomers**

**a – e** Representative micrographs at tilts of 0°, 20°, 30° and 40° with CHAPSO or LMNG (0.92 Å/pix) with corresponding 2D classes (scale bar, 180 Å). The untilted dataset with LMNG (**a**) displayed very strong particle orientation bias with only front views, while the untilted dataset with CHAPSO (**b**) contains side views at lower overall resolution of 2D class overviews. The dataset at 20° tilt (**c**) contains side views, the dataset at 30° tilt (**d**) contains side and top views, and the dataset at 40° tilt (**e**) is almost entirely composed of top views. The quality of the particles declined with higher stage tilts due to an increase in ice thickness, larger beam-induced movement, and poorer CTF estimation. This was evident by the resolution of the 2D classes and the declining CTF fit of the particle sets. The CTF was fitted to a range of 2.0–4.8 Å at 0° tilt, with the maximum fit increasing to 6.25 Å at 20° tilt, 7.9 Å at 30° tilt and 8.4 Å at 40° tilt. Hence, the tilt angle was limited to 40°.

**f** Flowchart of SPA processing. Map isotropy was improved by 2D class rebalancing, orientation rebalancing and limitation (cut-off filtering) of map amplitudes from overrepresented orientations.

**g** 2D classes (scale bar, 90 Å) of Mpn444 particles from the final map. Colored boxes indicate viewing direction (red for front views, blue for side views and yellow for top views). Viewing directions relative to the density map are displayed as orthogonal slices.

**h** Orientation diagnostics of the final map from cryoSPARC^42^. The diagram displays the relative signal amount in relation to the viewing direction. While our final dataset still oversampled particle front and back views (red boxes), we achieved sufficient signal contribution from all projection orientations (SCF* 0.880).

**i** Postprocessed electron density map of the Mpn444 monomer at 3.74 Å (EMD-55147).

**j** FSC curve calculated according to the gold-standard criterion of FSC at 0.143 (orange horizontal line).

**Supplemental Figure 2: Single-particle cryo-EM of Mpn436 monomers**

**a – b** Representative micrographs at tilts of 0° and 30° with and without LMNG (0.92 Å/pix) with corresponding 2D classes (scale bar, 180 Å). The untilted dataset (**a**) displayed very strong particle orientation bias with only front views, while the dataset at 30° tilt (**b**) contains side views and some top views.

**c** Flowchart of SPA processing. Map isotropy was improved by 2D class rebalancing, orientation rebalancing and limitation (cut-off filtering) of map amplitudes from overrepresented orientations. A refined consensus map was combined with a focused map of the lower half to yield the final composite map.

**d** 2D classes (scale bar, 90 Å) of Mpn436 particles from the final map. Colored boxes indicate viewing direction (red for front views, blue for side views and yellow for top views). Viewing directions relative to the density map are displayed as orthogonal slices.

**e** Orientation diagnostics of the final map from cryoSPARC^42^. The diagram displays the relative signal amount in relation to the viewing direction. While our final dataset still oversampled particle front and back views (red boxes), we achieved sufficient signal contribution from all projection orientations (SCF* 0.967).

**f** Postprocessed electron density maps of the Mpn436 monomer at 3.65 Å (consensus map, EMD-55129) and 4.07 Å (focused map, EMD-55131), as well as the postprocessed composite map (EMD-55148).

**g** FSC curves of the consensus map (left) and the focused map (right) calculated according to the gold-standard criterion of FSC at 0.143 (orange horizontal line).

**Supplemental Figure 3: Flexible loops of Mpn444 and Mpn436**

**a, b** (Top) Density map of the **(a)** Mpn444 monomer (EMD-55147) and **(b)** Mpn436 monomer (EMD-55148) (transparent grey surface) superimposed with the ribbon model of **(a)** Mpn444 and **(b)** Mpn436. The structure of the flexible loops (red) stands out from the density map. It is important to note that the loop conformation shown here (as deposited in PDB: 9SRQ for Mpn444 and PDB: 9SRR for Mpn436) is only one possible conformation. (Bottom) Structure of **(a)** Mpn444 and **(b)** Mpn436 as a ribbon model colored in grey (structured residues, as deposited in PDB: 9SRS for Mpn444 and PDB: 9SRV for Mpn436) and red (flexible residues).

**c** Density map of the Mpn444 trimer (EMD-53144, transparent grey surface) superimposed with the ribbon model of the Mpn444 trimeric assembly (as deposited in PDB: 9SSC). The structure of the flexible loops (red) stands out from the density map on the top view (facing away from the membrane). The loop conformation shown here is only one possible conformation. The central trimer and lateral dimer interfaces contain additional densities that are unaccounted for through rigid body docking of the monomers.

**Supplemental Figure 4: Amino acid sequence conservation of Mpn444**

Sequence alignment of the partial molecular model of Mpn444 without flexible loops, with the full-length model and homologs from five other *Mycoplasma* species (*M. genitalium*, *M. bradburyae*, *M. gallisepticum, M. testudineum*, *M. amphoriforme*). The alignment was done with the structure-guided sequence alignment tool PROMALS3D[17]. Conservation values >5, the consensus amino acid (aa) sequence and secondary structure (ss, h for helices and e for beta-strands) are displayed. Flexible loops missing in the partial model are shown in red with the label ‘loop’. It is apparent that the position of flexible loops correlates with poorly conserved regions of the sequence. Moreover, the loop regions in Mpn444 are predominantly absent in the homolog sequences. The position of the PPIase and chaperone domains are indicated using purple and green boxes. The catalytic residues of the PPIase domain are labelled with red stars.

**Supplemental Figure 5: Amino acid sequence conservation of Mpn436**

Sequence alignment of the partial molecular model of Mpn436 without flexible loops, with the full-length model and homologs from five other *Mycoplasma* species (*M. genitalium*, *M. bradburyae*, *M. gallisepticum, M. testudineum*, *M. amphoriforme*). The alignment was done with the structure-guided sequence alignment tool PROMALS3D[17]. Conservation values >5, the consensus amino acid (aa) sequence and secondary structure (ss, h for helices and e for beta-strands) are displayed. Flexible loops missing in the partial model are shown in red with the label ‘loop’. It is apparent that the position of flexible loops correlates with poorly conserved regions of the sequence. Moreover, the loop regions in Mpn436 are predominantly absent in the homolog sequences. The position of the PPIase and chaperone domains are indicated using purple and green boxes. The catalytic residues of the PPIase domain are labelled with red stars.

**Supplemental Figure 6: AlphaFold predictions of the proteins that share structural similarity with Mpn444**

Structure predictions of uncharacterized lipoproteins (**a**) Mpn444, (**b**) Mpn436, (**c**) Mpn489 from *Mycoplasma pneumoniae*, (**d**) MG309, € MG307, and (**f**) MG338 from *M. genitalium*; three uncharacterized *M. gallisepticum* proteins encoded by (**g**) *HFMG01WIA_4215*, (**h**) *GCW_03170*, and (**i**) *GCW_03165;* two DUF3713 domain-containing proteins from *M. tullyi* encoded by (**j**) *H3143_02785* and (**k**) *H3143_02790*; and (**l**) an uncharacterized *M. amphoriforme* protein encoded by *MAMA39_01690*. In addition, other uncharacterized mycoplasma proteins from *M. parvum* (*PRV_02465*), *M. suis* (*MSU_0344* and *MSUIS_02940*), *Candidatus Mycoplasma haemominutum* (*MHM_01560*), *M. haemolamae* (*MHLP_00685*), and *M. wenyonii* (*WEN_00705*) displayed similarity to the Mpn444 structure. AlphaFold2 (for a, d – i) and AlphaFold3 (for b, c) were used. The color scheme represents the confidence level of the prediction from AlphaFold (dark blue for very high confidence, light blue for high confidence, yellow for low confidence, dark orange for very low confidence).

**Supplemental Figure 7: Structural comparison of the PPIase domains from *M. pneumoniae* Mpn444 and Mpn436, and human Pin1**

**a** The PPIase domain of Mpn444 (purple) from *M. pneumoniae*. The three catalytic residues of Pin1 (S282, H468 and C/D284) are conserved in Mpn444 as S308, H494 and D310.

**b** The PPIase domain of Mpn436 (light purple) from *M. pneumoniae*. The catalytic residues of Pin1 (S282, H468 and D284) are partially conserved in Mpn436 as S326, T530 (for Pin1, catalytic activity has been shown for a HtoN mutant) and D327. **c** The PPIase domain of human Pin1 (yellow, PDB: 4TNS). The Pin1 structure is bound to the inhibitor all-trans retinoic acid (in black, with a black mesh surface representation). The substrate binding loop of Pin1, which recognizes phosphorylated peptides (in turquoise), is absent in Mpn444 and Mpn346.

**Supplemental Figure 8: Mpn444-D310A exhibits reduced PPIase activity while maintaining structural integrity**

**a** Cryo-EM map of Mpn444-D310A at 3.5 Å. Sky blue surface of the front view and side view.

**b** PPIase domains of Mpn444 (left, purple, aa 279-352, 471-504, 230-237) and Mpn444-D310A variant (right, pink) with red stars indicating the catalytic residues that are conserved among other PPIases.

**c** Superimposition of Mpn444 and Mpn444-D310 PPIase domains.

**d** PPIase activity assay. The increase in fluorescence intensity is directly proportional to the catalyzed proline cis to trans isomerization reaction. From left to right: Pin1 as positive control (dark blue), Mpn444 (red, 1 µg/µl), Mpn444-D310A (green, 1 µg/µl) The assay was done in two biological replicates and 35 technical replicates.

**Supplemental Figure 9: Chaperone-binding assay suggests that Mpn444 and Mpn436 interact with unfolded proteins**

The selective binding of Mpn444 and Mpn436 to unfolded client proteins was evaluated by size-exclusion chromatography (SEC). The proteins were eluted in Superose 6 Increase 10/300 GL column. The protein composition of the eluted fractions was assessed by SDS-PAGE.

**a** Control of the elution profile of native ADH

**b** Control of the elution profile Mpn444 with a molecular weight of approximately 140 kDa. Mpn444 was eluted around 21 ml.

**c** Control of the elution profile Mpn436 with a molecular weight of approximately 140 kDa. Mpn436 was eluted around 19 ml.

**d** Control of the elution profile of denatured ADH

**e** Chromatogram of Mpn444 and denatured ADH. The elution peak shifted from 21 to 17 ml. SDS gel confirms co-elution of Mpn444 and denatured ADH.

**f** Chromatogram of Mpn436 and denatured ADH. The elution peak shifted from 19 to 17 ml. SDS gel confirms co-elution of Mpn436 and denatured ADH

**g** Control for nonspecific binding. Lysozyme was used as a negative control. Lysozyme and denatured ADH did not co-elute.

**Supplemental Figure 10: Processing pipeline of the Mpn444 trimer**

**a** Representative micrographs at 0.92 Å/pix with corresponding 2D classes (scale bar 180 Å).

**b** 2D classes of dimeric complexes (incomplete trimers) (scale bar 180 Å).

**c** *Ab initio* reconstruction of the Mpn444 trimer structure with C1 symmetry (dark grey, top) and C3 symmetry (light grey, middle). The superimposition of the *ab initio* reconstructions (bottom, C3 symmetry structure surface in mesh representation) shows that both have the same overall domain architecture. The density is weaker in one monomer in the C1 symmetry structure, which can be attributed to the persistent presence of monomers and dimers in the particle set after 2D classification.

**d** Processing pipeline of the Mpn444 trimer and FSC curve of the final reconstruction from cryoSPARC.

**e** Density map of the Mpn444 trimer at 5.4 Å, colored by local resolution, with the corresponding FSC curve from cryoSPARC.

**Supplemental Figure 11: Comparison of the experimental trimer structure with the AlphaFold prediction**

**a** Density map of the Mpn444 trimer (transparent grey surface) fitted with the density maps of the Mpn444 monomer (dark blue, gray, sea green).

**b** Isosurface representation of the AlphaFold prediction of the Mpn444 trimer (dark blue, gray, sea green).

**c** Density map of the Mpn444 trimer (transparent grey surface) fitted with the molecular models of the Mpn444 monomer (ribbon models in dark slate grey, sea green, and a rainbow gradient from the N- to C-terminus). The N-terminal lipidated cysteines mediating membrane anchoring are indicated with red arrows.

**d** Ribbon model of the AlphaFold prediction of the Mpn444 trimer (dark slate grey, sea green, and a rainbow gradient from the N- to C-terminus).

**Supplemental Figure 12: Positions of external crosslinks on the Mpn444 homotrimer**

**a** Mpn444 interacts with other *M. pneumoniae* proteins^14^: Mpn436, Mpn489, Mpn396 (secD), Mpn141 (mgpA/major adhesion complex), Mpn376, and Mpn400 (ibpM).

**b** Residues are colored according to the crosslinking partner: Lys412 and Lys1151 (cyan) interact with Mpn489; Lys176 and Lys306 (lime) interact with Mpn436; Lys887 and Lys1058 (magenta) interact with Mpn396 (SecD); and Lys490 (yellow) interacts with Mpn141 (mgpA/major adhesion complex), Mpn376, and Mpn400 (ibpM).

Supplemental Figure 13: Structural comparison of the SEC translocon from M. pneumoniae *and* Thermus thermophilus.

**a** Experimental structures of the proteins SecDF (left, PDB: 3AQP, light blue), SecA (middle, PDB: 2IPC, sea green) and SecYEG (right, PDB: 5AWW, blue) forming the SEC translocon in *T. thermophilus*.

**b** AlphaFold3 predictions of the proteins SecD, SecA and SecYEG forming the SEC translocon in *M. pneumoniae.* The color scheme represents the confidence level of the prediction from AlphaFold (dark blue for very high confidence, light blue for high confidence, yellow for low confidence, dark orange for very low confidence).

**c** AlphaFold3 predictions of the SEC translocon from *T. thermophilus* (left) and *M. pneumoniae.* The colors are as in (a) for *T. thermophilus* and as in Figure 4 for *M. pneumoniae*.

**Supplemental Figure 14: Sub-tomogram average showing co-localization of a trimeric density extracellularly associated with the ribosome**

**a** Superimposed cryo-ET density maps of membrane-bound ribosome (in green) and extracellular trimeric density (in grey). Both maps were created with the same particle set (Cryo-ET Data Portal, dataset ID: DS-10003), but one was aligned on the ribosome (green, mask including ribosome and membrane) and the other on the extracellular density (grey, mask including extracellular density and membrane). The green map was fitted with a ribbon model of the Mycoplasma ribosome (PDB: 7PAK) and the grey map was fitted with the ribbon model of the homotrimeric Mpn444 assembly.

**b** Cryo-ET density maps of extracellular trimeric density. Shown in top and side view (top) as well as inverted in top and bottom view (bottom). The density map is fitted with the ribbon model of the homotrimeric Mpn444 assembly.

