## Supplementary material for "Structures of the essential *Mycoplasma pneumoniae* lipoproteins Mpn444 and Mpn436 reveal a peptidyl-prolyl isomerase domain involved in extracellular protein folding": Keles_Manger_SI_biorxiv_2026.pdf

Supplemental Information

### Supplemental Table 1: Single-particle cryo-EM data collection and processing

| | Mpn444 Monomer **EMD-55147** | Mpn444 Trimer **EMD-53144** | Mpn444-D310A | Mpn436 Monomer **EMD-55148** |
|---|---|---|---|---|
| **Microscope** | FEI Titan Krios | FEI Titan Krios | FEI Titan Krios | FEI Titan Krios |
| **Detector** | Gatan K3 Summit | Gatan K3 Summit | Gatan K3 Summit | Gatan K3 Summit |
| **Acquisition software** | SerialEM v4.1.0beta | SerialEM v4.1.0beta | SerialEM v4.1.0beta | SerialEM v4.1.0beta |
| **Magnification** | 105,000x | 105,000x | 105,000x | 105,000x |
| **Voltage (kV)** | 300 | 300 | 300 | 300 |
| **Electron exposure ($e^-$/Å$^2$)** | 50 | 50 | 50 | 50 |
| **Defocus range (µm)** | -0.8 to -3.6 | -0.8 to -3.6 | -0.8 to -3.6 | -0.8 to -3.6 |
| **Pixel size (Å)** | 0.92 | 0.92 | 0.92 | 0.92 |
| **Symmetry imposed** | C1 | C3 | C1 | C1 |
| **Initial particle images** | 6,832,694 | 3,376,184 | 4,992,685 | 6,582,748 |
| **Final particle images** | 432,841 | 506,291 | 159,443 | 468,211 |
| **Map resolution (Å)** | 3.74 | 5.85 | 5.2 | 3.65 |
| **FSC threshold** | 0.143 | 0.143 | 0.143 | 0.143 |
| **Map resolution range (Å)** | 3.2 - 5.5 | 3.5 - 7.5 | 3.2-4.5 | 3.1 – 4.9 |
| **Number of frames** | 50 | 50 | 50 | 50 |
| **Micrographs used** | 7,811 | 7,811 | 4,034 | 6,726 |
| **Processing software** | cryoSPARC v4.5.3 | cryoSPARC v4.5.3 | cryoSPARC v4.7.1 | cryoSPARC v4.5.3 |
| **Motion correction** | cryoSPARC v4.5.3 | cryoSPARC v4.5.3 | cryoSPARC v4.7.1 | cryoSPARC v4.5.3 |
| **CTF estimation** | cryoSPARC v4.5.3 | cryoSPARC v4.5.3 | cryoSPARC v4.7.1 | cryoSPARC v4.5.3 |
| **Particle images after 2D classification** | 2,992,076 | 1,126,935 | 2,135,473 | 2,168,069 |
| **Map sharpening B factor** | 104.1 | 377.8 | 129.4 | 107.9 |

### Supplemental Table 2: Model building statistics

| | Mpn444 | | Mpn436 | |
|---|---|---|---|---|
| | full-length | partial | full-length | partial |
| **PDB ID** | **9SRQ*** | **9SRS** | **9SRR** | **9SRV** |
| **Model composition** | | | | |
| Non-hydrogen atoms | 10,080 | 7,931 | 9,610 | 7,508 |
| Protein residues | 1,294 | 994 | 1,217 | 940 |
| Ligands | 0 | 0 | 0 | 0 |
| Bond lengths | 0.74 | 1.03 | 0.73 | 0.74 |

| | | | | |
|---|---|---|---|---|
| Outliers | 0 | 7 | 0 | 0 |
| Bond angles | 1.29 | 1.80 | 1.29 | 1.32 |
| Outliers | 29 | 29 | 12 | 15 |
| Chirality Outliers | 0 | 0 | 0 | 0 |
| Planarity Outliers | 4 | 8 | 3 | 6 |
| **Validation** | | | | |
| Clashscore | 1 | 8 | 1 | 1 |
| Poor rotamers (%) | 1.2 | 1.5 | 1.2 | 1.7 |
| Ramachandran plot | | | | |
| Favored (%) | 94 | 93 | 94 | 93 |
| Allowed (%) | 6 | 7 | 6 | 6 |
| Disallowed (%) | 0.1 | 0.5 | 0.4 | 0.5 |
| **Atom Inclusion** | 0.75 | 0.85 | 0.68 | 0.86 |
| **Q-Score** | 0.37 | 0.40 | 0.35 | 0.41 |

*PDB: 9SSC contains the homotrimeric assembly of PDB: 9SRQ after rigid-body fitting into the trimer density map, thus the model statistics are the same.

**Supplemental Table 3: Conservation of modelled and unmodelled regions**

Statistical evaluation of a sequence alignment of Mpn444 or Mpn436 from *Mycoplasma pneumoniae* with homologs from five other *Mycoplasma* species (*M. genitalium*, *M. bradburyae*, *M. gallisepticum, M. testudineum*, *M. amphoriforme*) (Supplemental Figure 4, Supplemental Figure 5). Residues within the stable fold are significantly more conserved (12% identity, 31% conservation for Mpn444, and 9% identity, 25% conservation for Mpn436) than residues on flexible loops (1% identity, 2% conservation for Mpn444, and 1% identity, 4% conservation for Mpn436). 'Conserved' includes amino acids with a >5 conservation score. 'Identical' includes residues that are unchanged across all homologs (labelled with a (9)). The total residue count excludes the N-terminal signal peptide, which was not part of the structure.

| | Mpn444 | | Mpn436 | |
|---|---|---|---|---|
| Total residue count | 1297 | | 1190 | |
| Residues modelled | 1042 | 90.3% | 940 | 79.0% |
| Conserved count | 322 | | 236 | |
| Conserved % of residues present | 30.9% | | 25.1% | |
| Identical (9) count | 125 | | 84 | |
| Identical (%) | 12.0% | | 9.0% | |
| Residues unmodelled (loop) | 255 | 19,7% | 277 | 21.0% |
| Conserved count | 6 | | 11 | |
| Conserved % | 2.4% | | 4.0% | |
| Identical (9) count | 3 | | 3 | |
| Identical (%) | 1.2% | | 1.1% | |
| Chaperone domain | 159 | 12.3% | 185 | 15.6% |
| Conserved count | 49 | | 47 | |
| Conserved % | 30.8% | | 25.4% | |
| Identical (9) count | 20 | | 19 | |
| Identical (%) | 12.5% | | 10.2% | |
| PPIase domain | 91 | 7.0% | 109 | 9.1% |
| Conserved count | 65 | | 47 | |
| Conserved % of PPIase domain | 71.4% | | 43.1% | |
| Identical (9) count | 32 | | 16 | |
| Identical (%) | 35.2% | | 14.7% | |

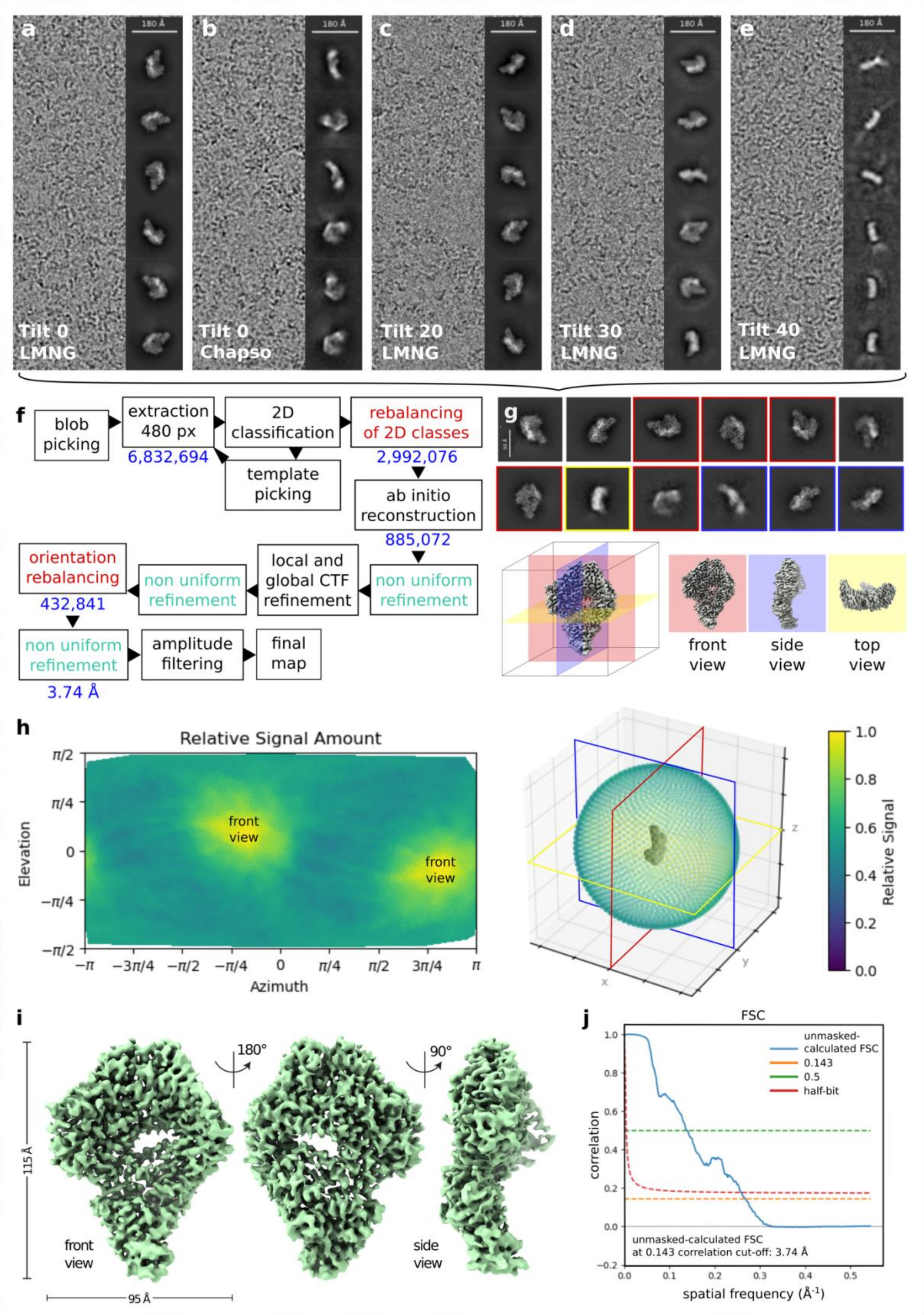

**Supplemental Figure 1: Single-particle cryo-EM of Mpn444 monomers**

**a – e** Representative micrographs at tilts of 0°, 20°, 30° and 40° with CHAPSO or LMNG (0.92 Å/pix) with corresponding 2D classes (scale bar, 180 Å). The untilted dataset with LMNG (**a**) displayed very strong particle orientation bias with only front views, while the untilted dataset with CHAPSO (**b**) contains side views at lower overall resolution of 2D class overviews. The dataset at 20° tilt (**c**) contains side views, the dataset at 30° tilt (**d**) contains side and top views, and the dataset at 40° tilt (**e**) is almost entirely composed of top views. The quality of the particles declined with higher stage tilts due to an increase in ice thickness, larger beam-induced movement, and poorer CTF estimation. This was evident by the resolution of the 2D classes and the declining CTF fit of the particle sets. The CTF was fitted to a range of 2.0–4.8 Å at 0° tilt, with the maximum fit increasing to 6.25 Å at 20° tilt, 7.9 Å at 30° tilt and 8.4 Å at 40° tilt. Hence, the tilt angle was limited to 40°.

**f** Flowchart of SPA processing. Map isotropy was improved by 2D class rebalancing, orientation rebalancing and limitation (cut-off filtering) of map amplitudes from overrepresented orientations.

**g** 2D classes (scale bar, 90 Å) of Mpn444 particles from the final map. Colored boxes indicate viewing direction (red for front views, blue for side views and yellow for top views). Viewing directions relative to the density map are displayed as orthogonal slices.

**h** Orientation diagnostics of the final map from cryoSPARC[42]. The diagram displays the relative signal amount in relation to the viewing direction. While our final dataset still oversampled particle front and back views (red boxes), we achieved sufficient signal contribution from all projection orientations (SCF* 0.880).

**i** Postprocessed electron density map of the Mpn444 monomer at 3.74 Å (EMD-55147).

**j** FSC curve calculated according to the gold-standard criterion of FSC at 0.143 (orange horizontal line).

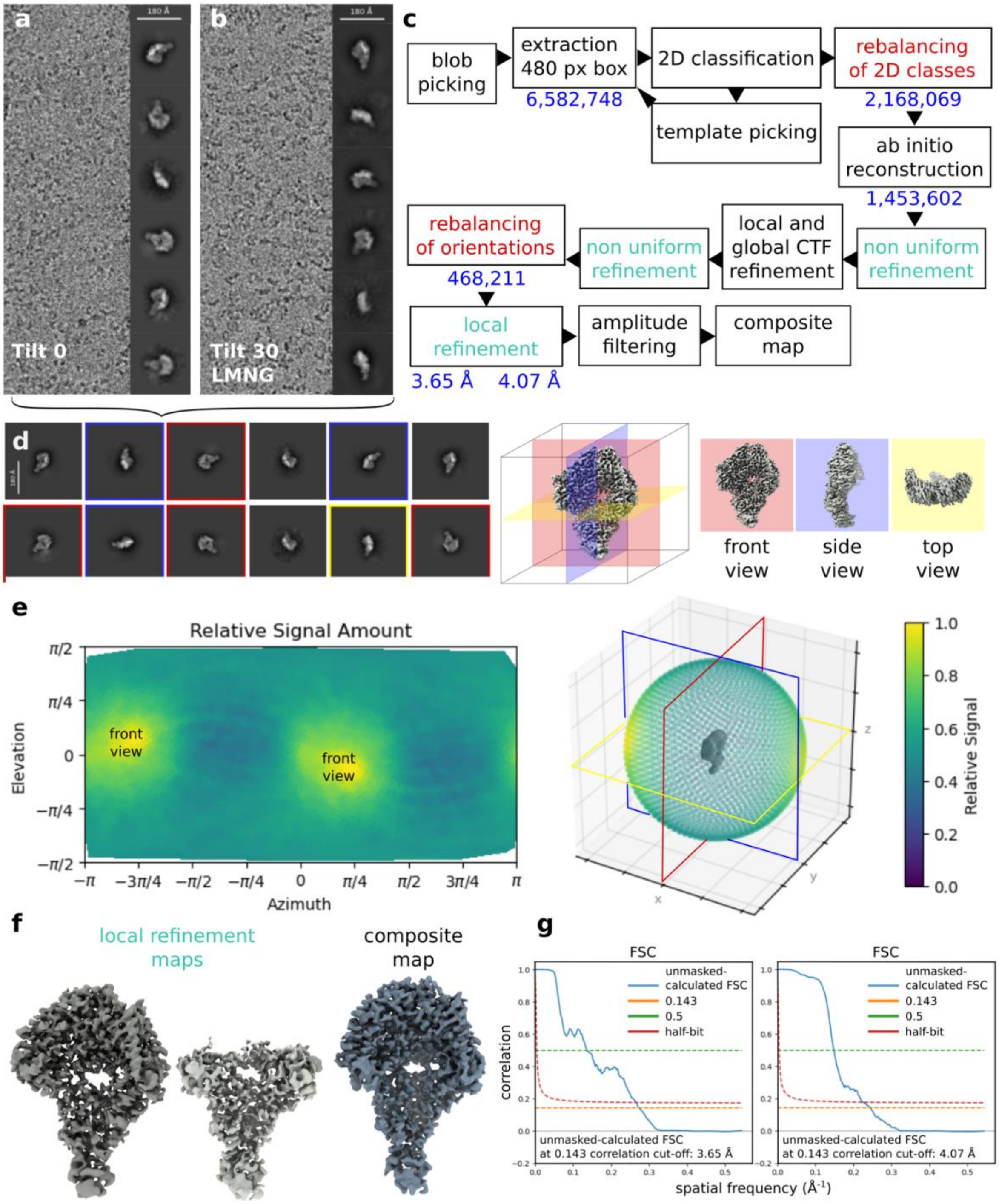

**Supplemental Figure 2: Single-particle cryo-EM of Mpn436 monomers**

**a – b** Representative micrographs at tilts of 0° and 30° with and without LMNG (0.92 Å/pix) with corresponding 2D classes (scale bar, 180 Å). The untilted dataset (**a**) displayed very strong particle orientation bias with only front views, while the dataset at 30° tilt (**b**) contains side views and some top views.

**c** Flowchart of SPA processing. Map isotropy was improved by 2D class rebalancing, orientation rebalancing and limitation (cut-off filtering) of map amplitudes from overrepresented orientations. A refined consensus map was combined with a focused map of the lower half to yield the final composite map.

**d** 2D classes (scale bar, 90 Å) of Mpn436 particles from the final map. Colored boxes indicate viewing direction (red for front views, blue for side views and yellow for top views). Viewing directions relative to the density map are displayed as orthogonal slices.
**e** Orientation diagnostics of the final map from cryoSPARC[42]. The diagram displays the relative signal amount in relation to the viewing direction. While our final dataset still oversampled particle front and back views (red boxes), we achieved sufficient signal contribution from all projection orientations (SCF* 0.967).
**f** Postprocessed electron density maps of the Mpn436 monomer at 3.65 Å (consensus map, EMD-55129) and 4.07 Å (focused map, EMD-55131), as well as the postprocessed composite map (EMD-55148).
**g** FSC curves of the consensus map (left) and the focused map (right) calculated according to the gold-standard criterion of FSC at 0.143 (orange horizontal line).

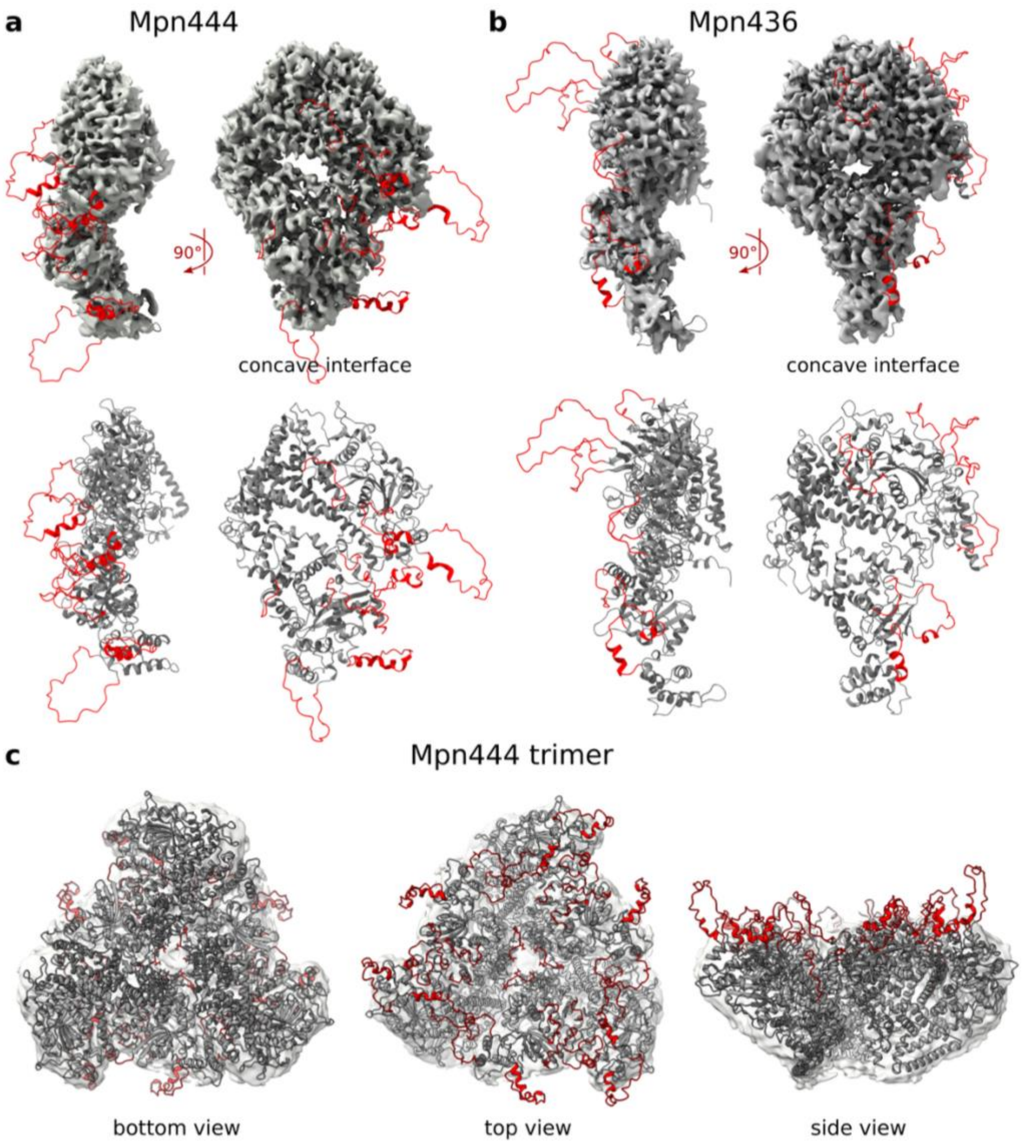

**Supplemental Figure 3: Flexible loops of Mpn444 and Mpn436**

**a, b** (Top) Density map of the **(a)** Mpn444 monomer (EMD-55147) and **(b)** Mpn436 monomer (EMD-55148) (transparent grey surface) superimposed with the ribbon model of **(a)** Mpn444 and **(b)** Mpn436. The structure of the flexible loops (red) stands out from the density map. It is important to note that the loop conformation shown here (as deposited in PDB: 9SRQ for Mpn444 and PDB: 9SRR for Mpn436) is only one possible conformation. (Bottom) Structure of **(a)** Mpn444 and **(b)** Mpn436 as a ribbon model colored in grey (structured residues, as deposited in PDB: 9SRS for Mpn444 and PDB: 9SRV for Mpn436) and red (flexible residues).

**c** Density map of the Mpn444 trimer (EMD-53144, transparent grey surface) superimposed with the ribbon model of the Mpn444 trimeric assembly (as deposited in PDB: 9SSC). The structure of the flexible loops (red) stands out from the density map on the top view (facing away from the membrane). The loop conformation shown here is only one possible conformation. The central trimer and lateral dimer interfaces contain additional densities that are unaccounted for through rigid body docking of the monomers.

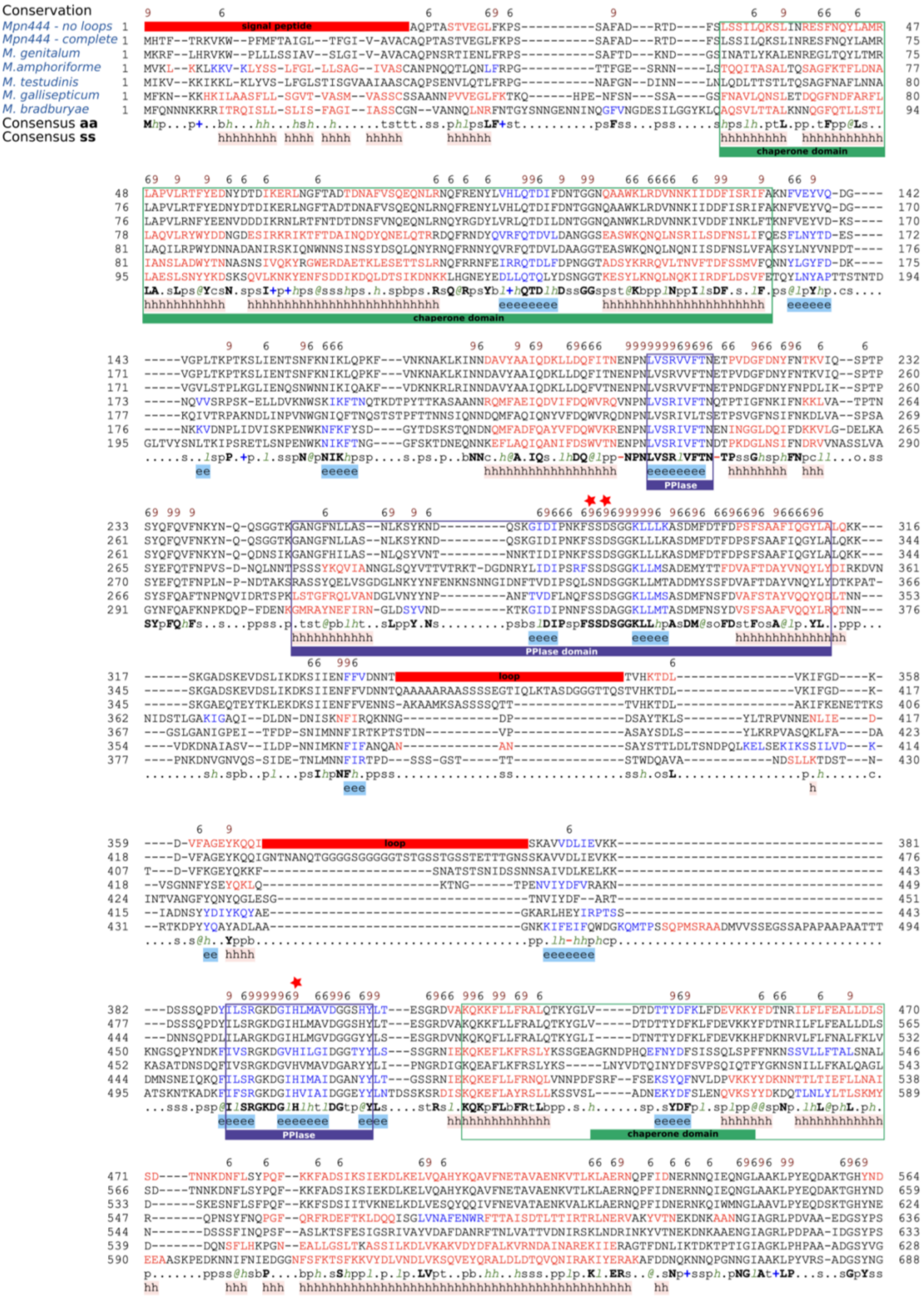
Conservation
Mpn444 - no loops
Mpn444 - complete
M. genitalum
M.amphoriforme
M. testudinis
M. gallisepticum
M. bradburyae
Consensus aa
Consensus ss
signal peptide
chaperone domain
PPIase
PPIase domain
loop

```
        6 9 6                                   6                    9 6        99                    69  99666
 565   LGNYYKDIIDNVDKKG     loop     NFSEEVVSKLKDNKKKVEEAAKKHVEALKVFTIPSPLYSQVILVQTK----LSFTPESTSLGLNLALN   644
 660   LGNYYKDIIDNVDKKGTSTVKTTSS-NTGQTKNFSEEVVSKLKDNKKKVEEAAKKHVEALKVFTIPSPLYSQVILVQTK----LSFTPESTSLGLNLALN   754
 625   LGIYYKDIIDKVSSNTSNSNSNSSS--SSSD-PFSKKIIDKLKENKKKVEAAVKKHVDELKVSVIPSPQYSQIILVDTK----LSSDPRNTSLALNLALN   717
 637   QEIYQLLVLNDWKNNQFKQSNV-LADDLSTK-GTKKYIFEQVKNAQENLQKEAIKLANALNLQSDSVSSFSQNIILRSN-------TDRFYSLPINLAIN   727
 634   QEIYELQLLKDFDP---KSTLT-IPANLSQT-GTRSFVLTQVQQTKAALDQAVTNLTTALNLRSDSVSQFSQNVFLASD-------TNSAYSLAINLAIN   721
 629   IESYYKNLLVENKL----LPENTLT-TKDQ--GLSKLFDQVVNDTKAKYFQTAKTVAEGLLLNGEPSYTYSQNVFVSSR-------YDPQYSLAANLAIN   714
 689   LEQYYLATFGLTGT------------PSTE--GTNKSIQDNVKKKREARDAKIQELLTKLKVETVLINSKSQNTFIDIKAEDEQFSKWENLKLALNLVLG   774
       b..Ybb.llsphp...............sp..shpcbl.pplpps+..hp..hpphhp.Lpl.s.sss.@SQslhlpoc.......psp.hSLtlNLAlN
       hhhhhhhhhhh                                   hhhhhhhhhhhhhhhhhh         eeeeeee             hhhhhhhhhhhh

         6     69  6  6      96   6             6                              6    6  6  6   9
 645   NYLTSTELQNSIKLSYFQEDEAFKKIIDITN-------LTFSQQSGGTGGTNGNNNL--TADN-WKIFKETYLLDLFESQAQKSIFGHVG   loop    724
 755   NYLTSTELQNSIKLSYFQEDEAFKKIIDITN-------LTFSQQSGGTGGTNGNNNL--TADN-WKIFKETYLLDLFESQAQKSIFGHVGSSDKNSSTKT   844
 718   AVLSSDELQNTIRRDYFVNDDQFKQAIDLDK-------LTFKNW----------NSL--NNEN-WNIFKYTYLFDLFQKQANPSIFGN-GVNESSTDNKP   796
 728   STIAAADTTNLVKNSFFTNSNDFKSFFDLTPTTN-SVYGTFKPTP---------TISKNVTDLLPETIRTVYGQTLWQALNDKSSYGVP-------ADIT   810
 722   ATITSTELTNNVKLSFFNNNKKFTDFFDYKANPNGQVYGTFKPTIFNTD----GDLSAVIND-LPNVIRTYYAQPMWQALGDKSGFGVPDIGDPT-KSLS   815
 715   TTIALPTSTNDFKRQFYLNDQGFTNFYDPRT-------NQFKDF----------EGL--KADQIKNVVNFYFIQSTWESQANKVKYGSW-------ATKA   788
 775   SIISSSLLTNEIKTEFIKSSKEFKEFYKVED-------GSVWSS----------FGI--DQTTLKNVIRNVYHFSVFSSLSDKTSKGVY-------TSFS   848
       shlstsphpNplK.p@@.psp.FpphhDhps........oFp............p.....s-...plh+.hYhbsh@ptbtpKo.@G.........ssbs
       hhhh  hhhhhhhhhhhh  hhhh                                   hhhhhhhhhhhhhhhhhhhh      eee

          6 6                      6     9   669  696 699 6 9           6   6   966
 725   ▮IEGVLDTLYSSLNLEERLD-----SDDVIDYLSYLYTAHWLLKDNLKNYKQSLQSKLSRTSNAFLVWSVDSEKNK          loop           794
 845   GIEGVLDTLYSSLNLEERLD-----SDDVIDYLSYLYTAHWLLKDNLKNYKQSLQSKLSRTSNAFLVWSVDSEKNKNDSQSTLSSTASSTSNTGLIQLRS   939
 797   KINGVLDSLYNSLNLEERLD-----SNDLINYYSYLYTVQWLLKDNLKNLKQNLQAKLSRTTNSFLVWSLASDKDRNNTASQAMSVSSSKS--------V   883
 811   SFNQIIDRIFTTRKTT---------SDDTLNYNTLIYTIKWLLENNLKNFKAIMQSRIAIGRQALALWSVSSSADKKLDDTS-----------------   883
 816   SFNQIIDRIFANKNI--TGVNANYDSADAISYYTFIYTLEWLLENNLQNFKNIMQFRVPVGTPALTTWTVPFAVNATL---------------------   891
 789   EYDNVLNNYFDNQRTISNLN-----DDARLNYLRYLNTFRYLIQDNLAEFKRILSTQITRGVSANVSWTLTNGTKLDNNGNIAS---------------   867
 849   TLNGIVNNLWNDQRLDKNPS-----SELTFNYYKYLYTLQWLLRNNLQNFKQIAKTSIKPGEVAFATWSLPVNLNMTSST-------------------   923
       php.llspl@ssbph..p.s.....SsshlsYhp@lYThpWLLcsNLpN@Kp.hpspls.spsAhhhWolss..s.s......................
       hhhhhhhhhhhhhh           hhhhhhhhhhhhhhhhhhh  hhhhhhhhhhh      eeeeeeee

                                996   99   66 9        6    6          66
 795               DNNSDITQTEVKNPNFVFGSSVYD                          loop                              818
 940   VVSLAQNQAAGQGGDNNSDITQTEVKNPNFVFGSSVYDWTNS------KTPEVNRAADDTSSFFYTKSSSSSTGAAQSSATVLRSLNQASGMTTKTAKN- 1032
 884   LVKMANNVAS----QTNQDFTKQEQQNPNYVFGSSAYNWTNN------KTPTVNSAANDISSLYYTKNNGSSSTSL--------TLMQKSAQQTNNQQR-  964
 884   --------------NGSNNPLTQYAANPNYLWGST-VNWQNHK----NAPSMADRGSDSKNPYQYTNLNN--------------------------NTK-  937
 892   --------------TAANNPLQTFANNPNYLWGSS-YNWQNHP----NAIANGDLGNTLANPYQYTSIDG--------------------------TSN-  945
 868   --------------TQDKINFTNFLANPNYLQGSK-FNLANL-----NKATDNNSNSQ-ANNLNYSTTNN--------------------------NQT-  919
 924   --------------DTNNNPLTKFDANPDGILGSKGSNWKNIVSSEASKIAEPNYWNNSSLTYKVSASED--------------------------SPKS  983
       ..............sssps.hppb..NPN@lhGSp.hsWpN........hspss.sss..sshbYop.ss..........................spp.
                          eeeee      ee                            eee

         699 9                  6   99   6                                          9 66 6 66    6   6  6    6
 819   ▮RYGFRGIVTSST--SGSLPEAVSRRLFKQFVNQ             loop                 TNNAYKGALFSF-GSMDNLKNIINGIQTQTE  879
1033   -RYGFRGIVTSST--SGSLPEAVSRRLFKQFVNQTEKGVKVGGQMLITTAKSGKATLVLKQQADDAESTTNNAYKGALFSF-GSMDNLKNIINGIQTQTE 1128
 965   -RFGFHGIVTNTS--SNNLPDAVRNRLFTSFVSQSEKSSSNGGQAQLQSTQSSGS--------------NETIYKGALFSF-GSLTKLIETIDNIPTQAE 1046
 938   -RYGFEGVKLRGD---TLNYSELRETLFDRYGSV-KD-------------------------------DKNVATGALYPYGNSRQTVVEYVQNLSFLSE 1000
 946   -RFGFNGVKLSGD---TLADTNLNTPLFSGYAQS-GN-------------------------------------QGVLYPYGNSRDSVIAFVDGIESNTV 1003
 920   -VYGFNGLSIANANNNTNLNTEVRSVLFDNFEQS-G--------------------------------------QNGVLYAFGGSLEDVVNYVNTLTTNRE  980
 984   PQYGFAGLVFKDS--SAPIAEEVKSALFTTYSSS-G--------------------------------------EKGTFFGYG-NKDNLIAYVDSIKSERE 1042
       .p@GFpGl.hpss...s.hsp.lpp.LFpp@spp.........................................pGhL@s@.sS.psl..hlsslpoppE
        eeeeeeeeee        hhhhhhhhhh                                               eeeee     hhhhhhhh    hhh

       66 6   6     6                          6                                      6 69 9969 66
 880   FDALYNHLTSDLNIDVTGVDK------NKTLTEQKTSLTSFVDSNFKQ             loop              KDVFSRFDGYIGDNKVEE  939
1129   FDALYNHLTSDLNIDVTGVDK------NKTLTEQKTSLTSFVDSNFKQSTQSAQRGDTSARSARSATVQIKKTQEDNQNTNYKDVFSRFDGYIGDNKVEE 1222
1047   FDALYNHLTSNLNINVTGVDR------SKSLQEQKTNLKNFANSNFNN----------------TQTVQLKQAQSKTNNSNFNDVFSRFEGYIGTNKT-- 1122
1001   LDGFIKFLASEAQIATNTIT-------GRNLNDKKAQTIALLNDT----------------------------------TKVNDGLFNRFSGYIGQNRS-- 1058
1004   LTNFVNVLATMGQVDTSSVPT------SNDLAARKTAVDALLRDV----------------------------------TKVPETLFTRFSGYIGTNKT-- 1062
 981   LDAFVELLRRAQV--NINLFETDANGMALSAEQRRANLISQLSDT----------------------------------KILPADVYTKFVGYVGQNKVSS 1045
1043   LDSLANRITDETRVPSNDYFKKDDKNNPLSYDKKLEEIKKMIN------------------------------------LIPASAFERFVGYIGNKKE-- 1104
       hDthhphLhs..plssssl.p......s.shppp+hpl.shlsss......................................slFsRFsGYIGpNKs..
       hhhhhhhhhhh                 hhhhhhhhhhhhhhh                                      hhhhhhhh      eeee

                           66 6966   66   69     66          69 96   669   66   9 6    9 69    66
 940   KN---YT-SYQFLSD--GGKYHATFVKQVNLDDVEKIG--TDSLKQEDSSKDKRLNLSLEEFLAAIALEALDPNNQTQAINALISGNK-------KGLVK 1024
1223   KN---YT-SYQFLSD--GGKYHATFVKQVNLDDVEKIG--TDSLKQEDSSKDKRLNLSLEEFLAAIALEALDPNNQTQAINALISGNK-------KGLVK 1307
1123   SN---YS-SYNFLQD--NQIYHAVYAKQINLEDVSMLG--SDSLNSTDSNNSKRLDLSLEEFLSTVALEALNPNNQTQAINALIANAK-------NGLVR 1207
1059   DK---LA-TLQAIRSPDQSKILPTFITQINYQDVNQLGQSSQWSANETSNNIYRLGLNLSEFLAIVVMQAMNADTQSNAITDLIRQNSNNQNNGTNGTVG 1154
1063   QD---IS-QLQAIRSSDGTHQLPTFVTQINHQDIQNIG-ASGWSLNDPANNKDRLGLNLDQFLAAIVMQAMDDGTQSNAISDLISSNVNPDN--SKGVIP 1155
1046   TDLTTQPDSFVPFTAAKSLLNWTTYIKQINFKEVANLG-GASWLT----NPSNRLDLSVDELLSIVAKYAFDNSYQNNAELAIISQQ---------SLID 1131
1105   EN---YDLN-NTLFVKGSLRRIATYVKQISYSDVDKLG--AGWLT----DSNKALGLSQSELLTIIAMTANNSAIQTQALTSLVNN----------KKLT 1184
       ps...hs.shphlps..s...hsT@lpQlNhpDVpplG..ss.......spppRLsLslpEhLthlhhpAhsss.QopAlssLIs.p.........thl.
       e       eeeeeee     eeeeeeeee hhhhhh                          hhhhhhhhhhhhhhh    hhhhhhhhhhh          ee

       6 9 96  66   66
1025   VGDFRIFSSISAQWVRRF--- 1042
1308   VGDFRIFSSISAQWVRRF--- 1325
1208   VGDNRLFSAISSQWVRKF--- 1225
1155   VGDRRLLNALTTRWAHSQSRN 1175
1156   IGDKRLLDALTTRWA--KALN 1174
1132   VNDKRLYDALGLRWVLRNS-- 1150
1185   VRDVRLFGALGSFFAKKIN-- 1203
       VsD.RlhstlsspWh.pb...
       e  ee      hhhhhh
```

**Supplemental Figure 4: Amino acid sequence conservation of Mpn444**

Sequence alignment of the partial molecular model of Mpn444 without flexible loops, with the full-length model and homologs from five other *Mycoplasma* species (*M. genitalium*, *M. bradburyae*, *M. gallisepticum, M. testudineum*, *M. amphoriforme*). The alignment was done with the structure-guided sequence alignment tool PROMALS3D[17]. Conservation values >5, the consensus amino acid (aa) sequence and secondary structure (ss, h for helices and e for beta-strands) are displayed. Flexible loops missing in the partial model are shown in red with the label ‘loop’. It is apparent that the position of flexible loops correlates with poorly conserved regions of the sequence. Moreover, the loop regions in Mpn444 are predominantly absent in the homolog sequences. The position of the PPIase and chaperone domains are indicated using purple and green boxes. The catalytic residues of the PPIase domain are labelled with red stars.

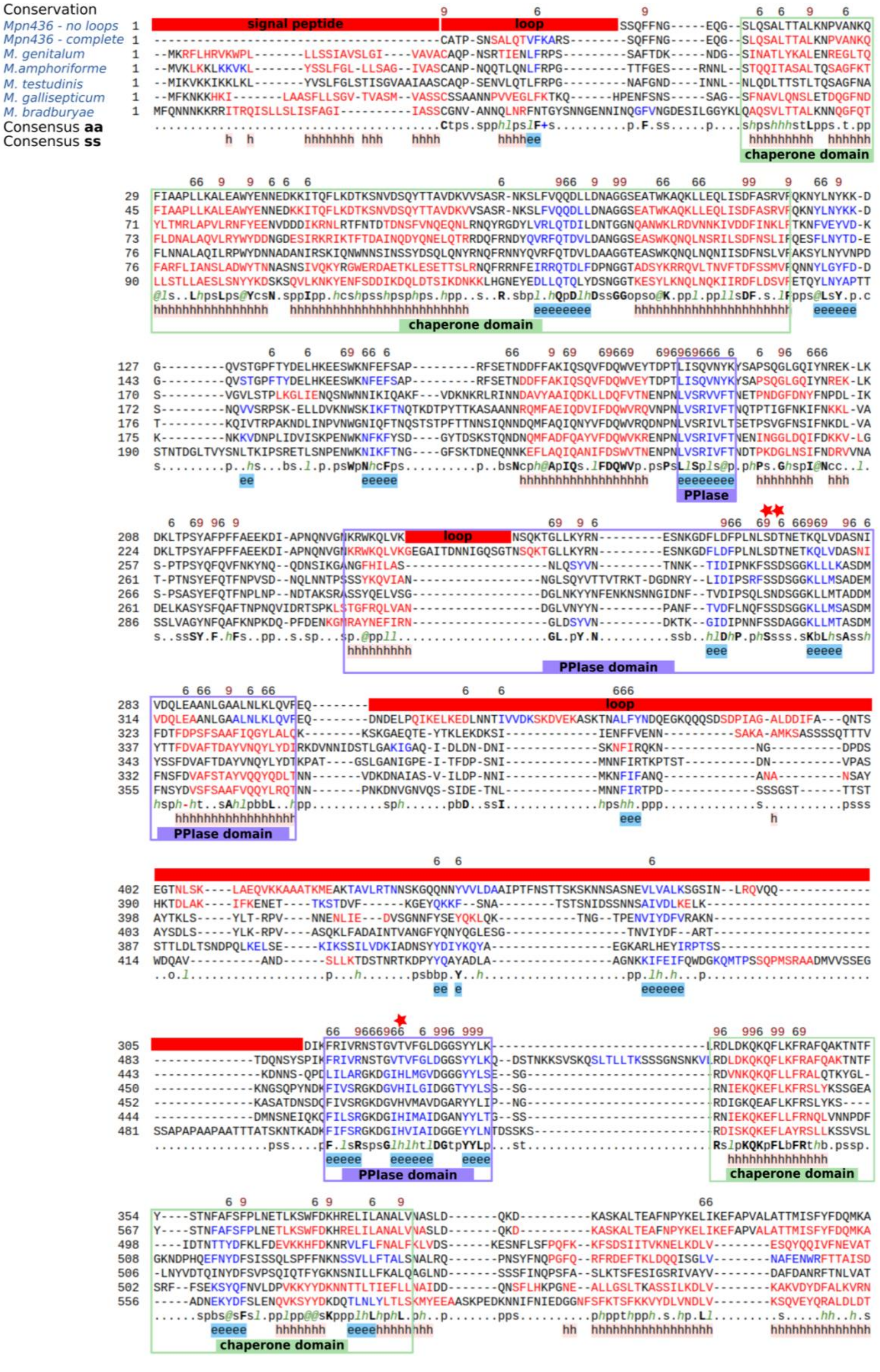
Conservation
Mpn436 - no loops
Mpn436 - complete
M. genitalum
M.amphoriforme
M. testudinis
M. gallisepticum
M. bradburyae
Consensus aa
Consensus ss
signal peptide
loop
chaperone domain
PPIase
PPIase domain

```
      6   6     6          69666 99       9 9      9
433   LNNKLLE  [loop]  NGLSAKLPYVN--TNGNYEKLNNYFTFLITKTLWPKVGQEETSISEESNKLKTKTADVDKIRDKILENIQTKV  510
646   LNNKLLERARNLNQNV--NQANPTPWLNGLSAKLPYVN--TNGNYEKLNNYFTFLITKTLWPKVGQEETSISEESNKLKTKTADVDKIRDKILENIQTKV  741
578   AENKVALKLAERNQPFIDNERNKQIWMNGLAAVLPYEQDSKTGHYNELGIYYKDIIDKV--SSNTSNSNSNSSSSS-SDPFSKKIIDKLKENKKKVEAAV  674
591   TLTTIRTRLNERVAKYVTNEKDNKAANNGIAGRLPDVAA-EDGSYPSQEIYQLLVLNDW--KNNQFKQSNVLADDLSTKGTKKYIFEQVKNAQENLQKEA  687
588   TVDNIRSKLNDRINKYVTNEKDNKAAENGIAGRLPDPAA-IDGSYPSQEIYELQLLKDF-----DPKSTLTIPANLSQTGTRSFVLTQVQQTKAALDQAV  681
583   DAINAREKIIERAGTFDNLIKTDKTPTIGIAGKLPHPAA-ADGSYVGIESYYKNLLVEN------KLLPENTLTTK-DQGLSKLFDQVVNDTKAKYFQTA  674
643   QVQNIRAKIYERAKAFDDNQKNNQPGNNGIAAKLPYVRS-ADGSYNGLEQYYLATFG------------LTGTPS-TEGTNKSIQDNVKKKREARDAKI  727
      ..ppl..+h.p.s..h..Np.ssps..NGltt+LP.s....sGsY..bp.Yb..llsch.......ppsp...ps...ps.p..l.p..pp.b..hp..h
      hhhhhhhhhhhhh                                   hhhhhhhh                                hhhhhhhhhh

      9            96                66  96 6   6          6         6   96 6 6            6
511   NDFVKNKLKPALAPRPAYSNVILLNVN-------NDKVLSSGANWSLASLLQSDKVNPLSFMLLKQAFDNNDLFKKAQKLFKD------IQEKSSN----  593
742   NDFVKNKLKPALAPRPAYSNVILLNVN-------NDKVLSSGANWSLASLLQSDKVNPLSFMLLKQAFDNNDLFKKAQKLFKD------IQEKSSN----  824
675   KKHV-DELKVSVIPSPQYSQIILVDTK----LSSDPRNTSLALNLALNAVLSS---DELQNTIRRDYFVNDDQFKQAIDLDK-------LTFKNWNS---  756
688   IKLA-NALNLQSDSVSSFSQNIILRSN-------TDRFYSLPINLAINSTIAA---ADTTNLVKNSFFTNSNDFKSFFDLTPTTN-SVYGTFKPT-----  770
682   TNLT-TALNLRSDSVSQFSQNVFLASD-------TNSAYSLAINLAINATITS---TELTNNVKLSFFNNNKKFTDFFDYKANPNGQVYGTFKPTIFNTD  770
675   KTVA-EGLLLNGEPSYTYSQNVFVSSR-------YDPQYSLAANLAINTTIAL---PTSTNDFKRQFYLNDQGFTNFYDPRT-------NQFKDFEG---  753
728   QELL-TKLKVETVLINSKSQNTFIDIKAEDEQFSKWENLKLALNLVLGSIISS---SLLTNEIKTEFIKSSKEFKEFYKVED-------GSVWSSFG---  813
      pphh.s.Lp....s.s.@Spslhlssp.......psp.hS.thNhtlsthl.t...s.ho..lbpph@.Nsp.Fpphbch..........pbKs......
      hhhh hhh           eeeeeee           hhhhhhhhhhhhhhh        hhhhhhhhhhh   hhhh

                           6    6  6 66   9             6                                6   669  69
594   ---NGGMQSSSTTN-SDADALSKVIGNYYYTTWAKLTDKSIYGNPK-------DNKFDELFKLAFEASIDEKSFNVD-----YKAVIDHYRFIYTLQWLV  677
825   ---NGGMQSSSTTN-SDADALSKVIGNYYYTTWAKLTDKSIYGNPK-------DNKFDELFKLAFEASIDEKSFNVD-----YKAVIDHYRFIYTLQWLV  908
757   ---------------LNNENWNIFKYTYLFDLFQKQANPSIFGNGVNESSTDNKPKINGVLDSLYNSLNLEE--RLD-----SNDLINYYSYLYTVQWLL  834
771   PTIS---------KNVTDLLPETIRTVYGQTLWQALNDKSSYGVPA------DITSFNQIIDRIFTTRKTT-----------SDDTLNYNTLIYTIKWLL  844
771   GDLS---------AVIND-LPNVIRTYYAQPMWQALGDKSGFGVPDIGDPTKSLSSFNQIIDRIFANKNI----TGVNANYDSADAISYYTFIYTLEWLL  856
754   -------------L-KADQIKNVVNFYFIQSTWESQANKVKYGSWA------TKAEYDNVLNNYFDNQRTIS--NLN-----DDARLNYLRYLNTFRYLI  826
814   -------------I-DQTTLKNVIRNVYHFSVFSSLSDKTSKGVYT------SFSTLNGIVNNLWNDQRLDK--NPS-----SELTFNYYKYLYTLQWLL  886
      ................ssph.s.l...Yhbsh@..bssKo.@Gs..........sphs.lhp.h@psp..p...s.s......pshls@hp@lYTlpWLl
                      hhhhhhhhhhhhhhhhh     eee             hhhhhhhhhhhhhh           hhhhhhhhhhhhhhhhhh

        69 669         6  6   6                                     99 6  99   9  9
678   DQKLKNFKSLLKTNLKFGEVAFIAYK-------------------------------------NTETTNFSNPQGVFGS  [loop]  KESTQ  724
909   DQKLKNFKSLLKTNLKFGEVAFIAYK-------------------------------------NTETTNFSNPQGVFGSY-FNYENSAS----EVKESTQ  966
835   KDNLKNLKQNLQAKLSRTTNSFLVWSLASDKDRNNTASQAMSVSSSKSVLVKMANNVASQTNQDFTKQEQQNPNYVFGSSAYNWTNNK------TPTVNS  928
845   ENNLKNFKAIMQSRIAIGRQALALWSVSSSADKKLDD--------------------TSNGSNNPLTQYAANPNYLWGST-VNWQNHKN----APSMADR  919
857   ENNLQNFKNIMQFRVPVGTPALTTWTVPFAVNATL------------------------TAANNPLQTFANNPNYLWGSS-YNWQNHPN----AIANGDL  927
827   QDNLAEFKRILSTQITRGVSANVSWTLTNGTKLDNNGN------------------IASTQDKINFTNFLANPNYLQGSK-FNLANLNK-----ATDNNS  902
887   RNNLQNFKQIAKTSIKPGEVAFATWSLPVNLNMTS---------------------STDTNNNPLTKFDANPDGILGSKGSNWKNIVSSEASKIAEPNY  964
      cppLpNFKplhphpl..GpsAhhh@p....................................ssbpp..sNPp.lhGS..hN@pN........h.pss.
      hh hhhhhhhhhhh      eeeeeee                                          eee          e

            6                        699 9                66
725   TLD-PNNFFYKTTTKP  [loop]  TDHYGFTGLSTSTSS---MFDASSRDAILQQI---------------------------T  769
967   TLD-PNNFFYKTTTKPTVQAIQQVASLALVQKQQMQQNSTDHYGFTGLSTSTSS---MFDASSRDAILQQI---------------------------T 1034
929   AANDISSLYYTKNNGS-----SSTSLTLMQKSAQQTNNQQRRFGFHGIVTNTSS--NNLPDAVRNRLFTSFVSQSEKSSSNGGQAQLQSTQSSGSNETIY 1021
920   GSDSKNPYQYTNLNN-----------------------NTKRYGFEGVKLRGDT---LNYSELRETLFDRYGSVKD------------------DKNVA  974
928   GNTLANPYQYTSIDG-----------------------TSNRFGFNGVKLSGDT---LADTNLNTPLFSGYAQSGN----------------------  977
903   NSQ-ANNLNYSTTNN-----------------------NQTVYGFNGLSIANANNNTNLNTEVRSVLFDNFEQSG----------------------Q  954
965   WNNSSLTYKVSASED---------------------SPKSPQYGFAGLVFKDSS--APIAEEVKSALFTTYSSSG----------------------E 1017
      s.s..sshbYps.s.......................pppp@GFpGl.hpsso....hssp.+sslhpph..............................
           eee                             eeeeeeeee         hhhhhhhhhh

       6 6  69 6   6   6        66      6
770   KTSLQQYG-SKDQLKKIIQGTNNQLLLDRIAVQLSGL-NPSTTNGGSGKTIATYFQVDAVGNPTLDFQAKRKLLL-DLL--------------------  845
1035  KTSLQQYG-SKDQLKKIIQGTNNQLLLDRIAVQLSGL-NPSTTNGGSGKTIATYFQVDAVGNPTLDFQAKRKLLL-DLL-------------------- 1110
1022  KGALFSFG-SLTKLIETIDNIPTQAEFDALYNHLTSN-LNINVTGVDRSK--------------SLQEQKTNLKNFANSNFNNTQTVQLKQAQSKTNNS 1104
975   TGALYPYGNSRQTVVEYVQNLSFLSELDGFIKFLASE-AQIATNTITGR---------------NLNDKKAQTIALLNDT-----------------TK 1040
978   QGVLYPYGNSRDSVIAFVDGIESNTVLTNFVNVLATM-GQVDTSSVPTSN--------------DLAARKTAVDALLRDV-----------------TK 1044
955   NGVLYAFGGSLEDVVNYVNTLTTNRELDAFVELLRRAQVNINLFE----------TDANGMAL-SAEQRRANLISQLSDT-----------------KI 1025
1018  KGTFFGYG-NKDNLIAYVDSIKSERELDSLANRITDE-TRVPSNDYFK--------KDDKNNPL-SYDKKLEEIKKMIN--------------------L 1086
      pssLb.@G.S.ppl..hlpshpsp.bLD.hh..Lss..s..shss.s...................shp.++..lb..l......................
       eeeee    hhhhhhhh    hhhhhhhhhhhhhh                              hhhhhhhhhhhhh

          6  6 96 6 6                          696   66  66                 6     6      9 6   9
846   ---DQYQNYFGN  [loop]  GTGN-YLTYQNGSDKY---TYTQFTYQDIDSLS--LTTT----SGTNN--KIASDVVAALLLFQAADKGTQQL  915
1111  ---DQYQNYFGNGAQKSQRD---STPSGTGN-YLTYQNGSDKY---TYTQFTYQDIDSLS--LTTT----SGTNN--KIASDVVAALLLFQAADKGTQQL 1192
1105  NFNDVFSRFEGYIGTNKTSNYS--------SYNFLQD--NQIYHAVYAKQINLEDVSMLG--SDSLNSTDSNNSKRLDLSLEEFLSTVALEALNPNNQTQ 1192
1041  VNDGLFNRFSGYIGQNRSDKLA--------TLQAIRSPDQSKILPTFITQINYQDVNQLGQSSQWSANETSNNIYRLGLNLSEFLAIVVMQAMNADTQSN 1132
1045  VPETLFTRFSGYIGTNKTQDIS--------QLQAIRSSDGTHQLPTFVTQINHQDIQNIG-ASGWSLNDPANNKDRLGLNLDQFLAAIVMQAMDDGTQSN 1135
1026  LPADVYTKFVGYVGQNKVSSTDLTTQP--DSFVPFTAAKSLLNWTTYIKQINFKEVANLG-GASWL----TNPSNRLDLSVDELLSIVAKYAFDNSYQNN 1118
1087  IPASAFERFVGYIGNKKEENYD--------LNNTLFVKGSLRRIATYVKQISYSDVDKLG--AGWL----TDSNKALGLSQSELLTIIAMTANNSAIQTQ 1172
      ...s.@pp@.G..tppppps..........p..hh..sss......hhpQhs@pDlspLt...sh.....osspp...ls.s.hhthlhhpAhs.shQp.
       hhhhhhhh      eee              eeee       eeeeeeee hhh                    hhhhhhhhhhhhhhh    hhhh

      9   6                   6 9 96  66
916   ALSAINKP-----------QLNIGDKRIESGLKL---LK----   940
1193  ALSAINKP-----------QLNIGDKRIESGLKL---LK----  1217
1193  AINALIANAK-------NGLVRVGDNRLFSAISSQWVRKF---  1225
1133  AITDLIRQNSNNQNNGTNGTVGVGDRRLLNALTTRWAHSQSRN  1175
1136  AISDLISSNVNPDN--SKGVIPIGDKRLLDALTTRWA--KALN  1174
1119  AELAIISQQ----------SLIDVNDKRLYDALGLRWVLRNS--  1150
1173  ALTSLVNN-----------KKLTVRDVRLFGALGSFFAKKIN--  1203
      Alssl.p................lslsDpRlbstL......+....
      hhhhhhh                eee  ee hhh  hhhhh
```

**Supplemental Figure 5: Amino acid sequence conservation of Mpn436**

Sequence alignment of the partial molecular model of Mpn436 without flexible loops, with the full-length model and homologs from five other *Mycoplasma* species (*M. genitalium*, *M. bradburyae*, *M. gallisepticum, M. testudineum*, *M. amphoriforme*). The alignment was done with the structure-guided sequence alignment tool PROMALS3D[17]. Conservation values >5, the consensus amino acid (aa) sequence and secondary structure (ss, h for helices and e for beta-strands) are displayed. Flexible loops missing in the partial model are shown in red with the label 'loop'. It is apparent that the position of flexible loops correlates with poorly conserved regions of the sequence. Moreover, the loop regions in Mpn436 are predominantly absent in the homolog sequences. The position of the PPIase and chaperone domains are indicated using purple and green boxes. The catalytic residues of the PPIase domain are labelled with red stars.

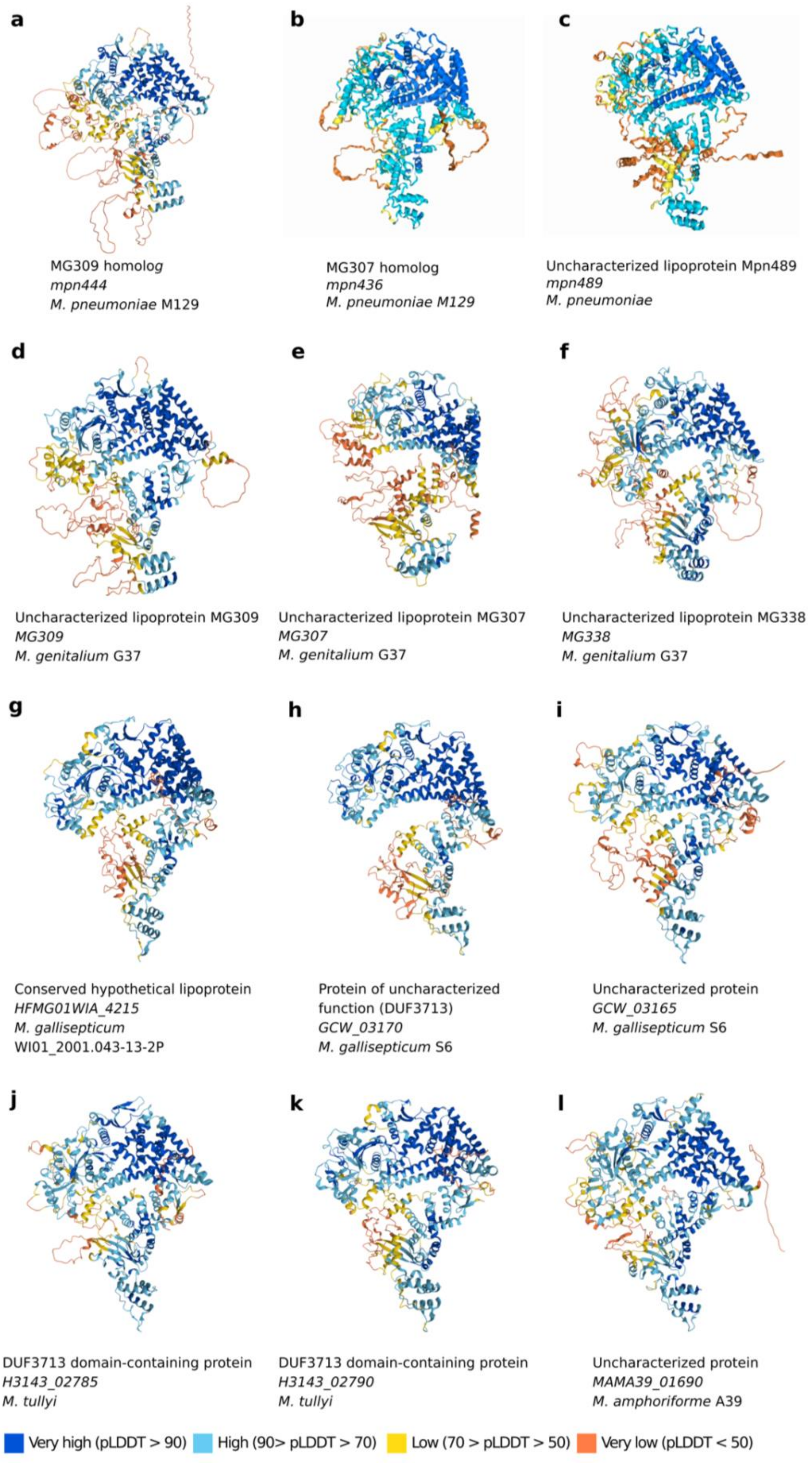
a
MG309 homolog
mpn444
M. pneumoniae M129
b
MG307 homolog
mpn436
M. pneumoniae M129
c
Uncharacterized lipoprotein Mpn489
mpn489
M. pneumoniae
d
Uncharacterized lipoprotein MG309
MG309
M. genitalium G37
e
Uncharacterized lipoprotein MG307
MG307
M. genitalium G37
f
Uncharacterized lipoprotein MG338
MG338
M. genitalium G37
g
Conserved hypothetical lipoprotein
HFMG01WIA_4215
M. gallisepticum
WI01_2001.043-13-2P
h
Protein of uncharacterized function (DUF3713)
GCW_03170
M. gallisepticum S6
i
Uncharacterized protein
GCW_03165
M. gallisepticum S6
j
DUF3713 domain-containing protein
H3143_02785
M. tullyi
k
DUF3713 domain-containing protein
H3143_02790
M. tullyi
l
Uncharacterized protein
MAMA39_01690
M. amphoriforme A39
Very high (pLDDT > 90)
High (90> pLDDT > 70)
Low (70 > pLDDT > 50)
Very low (pLDDT < 50)

**Supplemental Figure 6: AlphaFold predictions of the proteins that share structural similarity with Mpn444**

Structure predictions of uncharacterized lipoproteins (**a**) Mpn444, (**b**) Mpn436, (**c**) Mpn489 from *Mycoplasma pneumoniae*, (**d**) MG309, € MG307, and (**f**) MG338 from *M. genitalium*; three uncharacterized *M. gallisepticum* proteins encoded by (**g**) *HFMG01WIA_4215*, (**h**) *GCW_03170*, and (**i**) *GCW_03165;* two DUF3713 domain-containing proteins from *M. tullyi* encoded by (**j**) *H3143_02785* and (**k**) *H3143_02790* ; and (**l**) an uncharacterized *M. amphoriforme* protein encoded by *MAMA39_01690*. In addition, other uncharacterized mycoplasma proteins from *M. parvum* (*PRV_02465*), *M. suis* (*MSU_0344* and *MSUIS_02940*), *Candidatus Mycoplasma haemominutum* (*MHM_01560*), *M. haemolamae* (*MHLP_00685*), and *M. wenyonii* (*WEN_00705*) displayed similarity to the Mpn444 structure. AlphaFold2 (for a, d – i) and AlphaFold3 (for b, c) were used. The color scheme represents the confidence level of the prediction from AlphaFold (dark blue for very high confidence, light blue for high confidence, yellow for low confidence, dark orange for very low confidence).

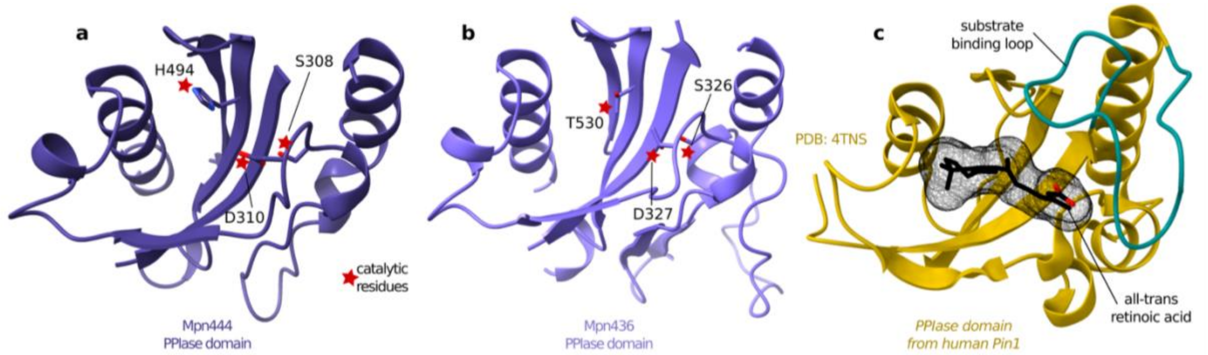

**Supplemental Figure 7: Structural comparison of the PPIase domains from *M. pneumoniae* Mpn444 and Mpn436, and human Pin1**

**a** The PPIase domain of Mpn444 (purple) from *M. pneumoniae*. The three catalytic residues of Pin1 (S282, H468 and C/D284) are conserved in Mpn444 as S308, H494 and D310.

**b** The PPIase domain of Mpn436 (light purple) from *M. pneumoniae*. The catalytic residues of Pin1 (S282, H468 and D284) are partially conserved in Mpn436 as S326, T530 (for Pin1, catalytic activity has been shown for a HtoN mutant) and D327.

**c** The PPIase domain of human Pin1 (yellow, PDB: 4TNS). The Pin1 structure is bound to the inhibitor all-trans retinoic acid (in black, with a black mesh surface representation). The substrate binding loop of Pin1, which recognizes phosphorylated peptides (in turquoise), is absent in Mpn444 and Mpn346.

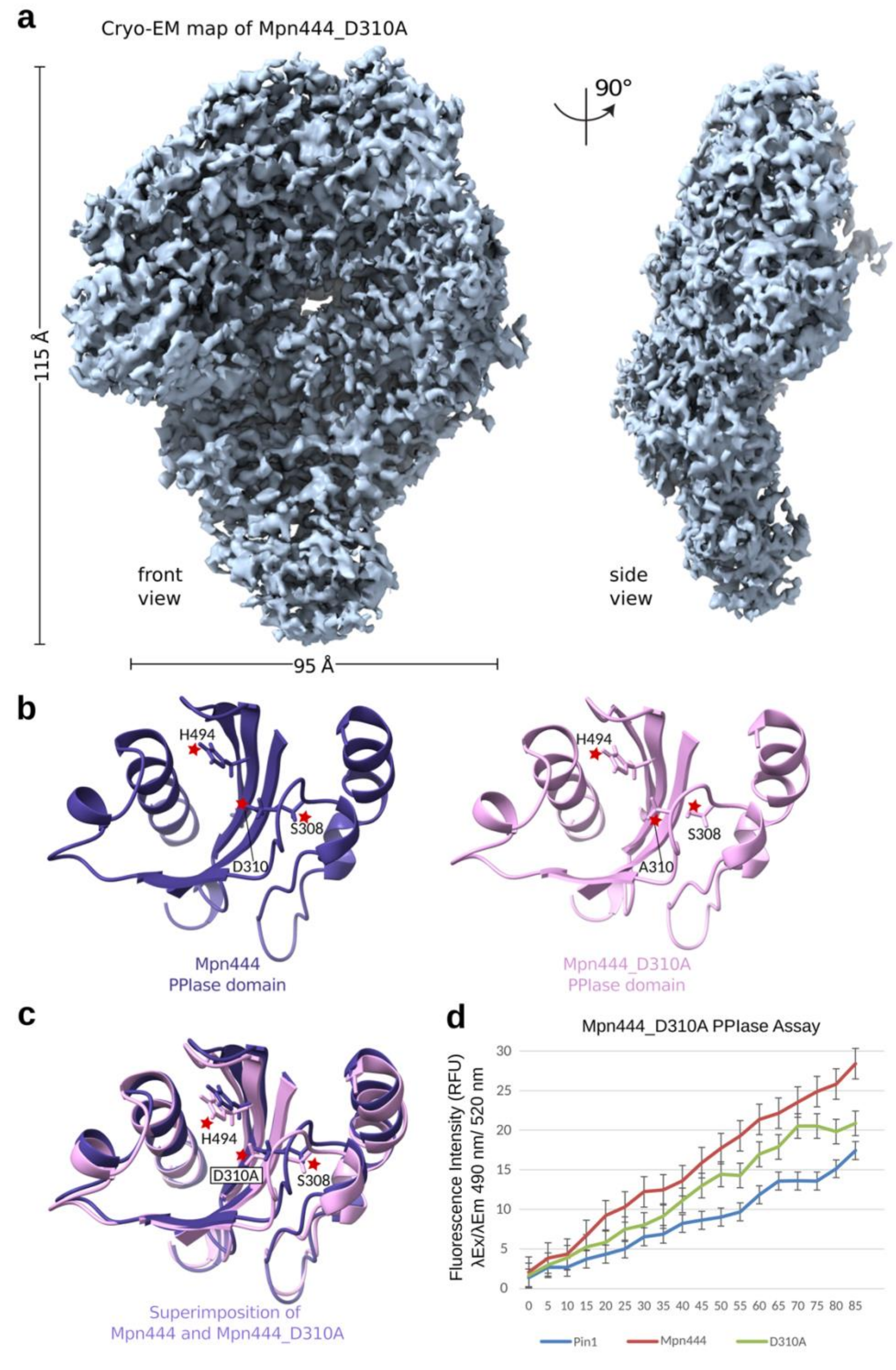

**Supplemental Figure 8: Mpn444-D310A exhibits reduced PPIase activity while maintaining structural integrity**

**a** Cryo-EM map of Mpn444-D310A at 3.5 Å. Sky blue surface of the front view and side view.

**b** PPIase domains of Mpn444 (left, purple, aa 279-352, 471-504, 230-237) and Mpn444-D310A variant (right, pink) with red stars indicating the catalytic residues that are conserved among other PPIases.
**c** Superimposition of Mpn444 and Mpn444-D310 PPIase domains.
**d** PPIase activity assay. The increase in fluorescence intensity is directly proportional to the catalyzed proline cis to trans isomerization reaction. From left to right: Pin1 as positive control (dark blue), Mpn444 (red, 1 µg/µl), Mpn444-D310A (green, 1 µg/µl) The assay was done in two biological replicates and 35 technical replicates.

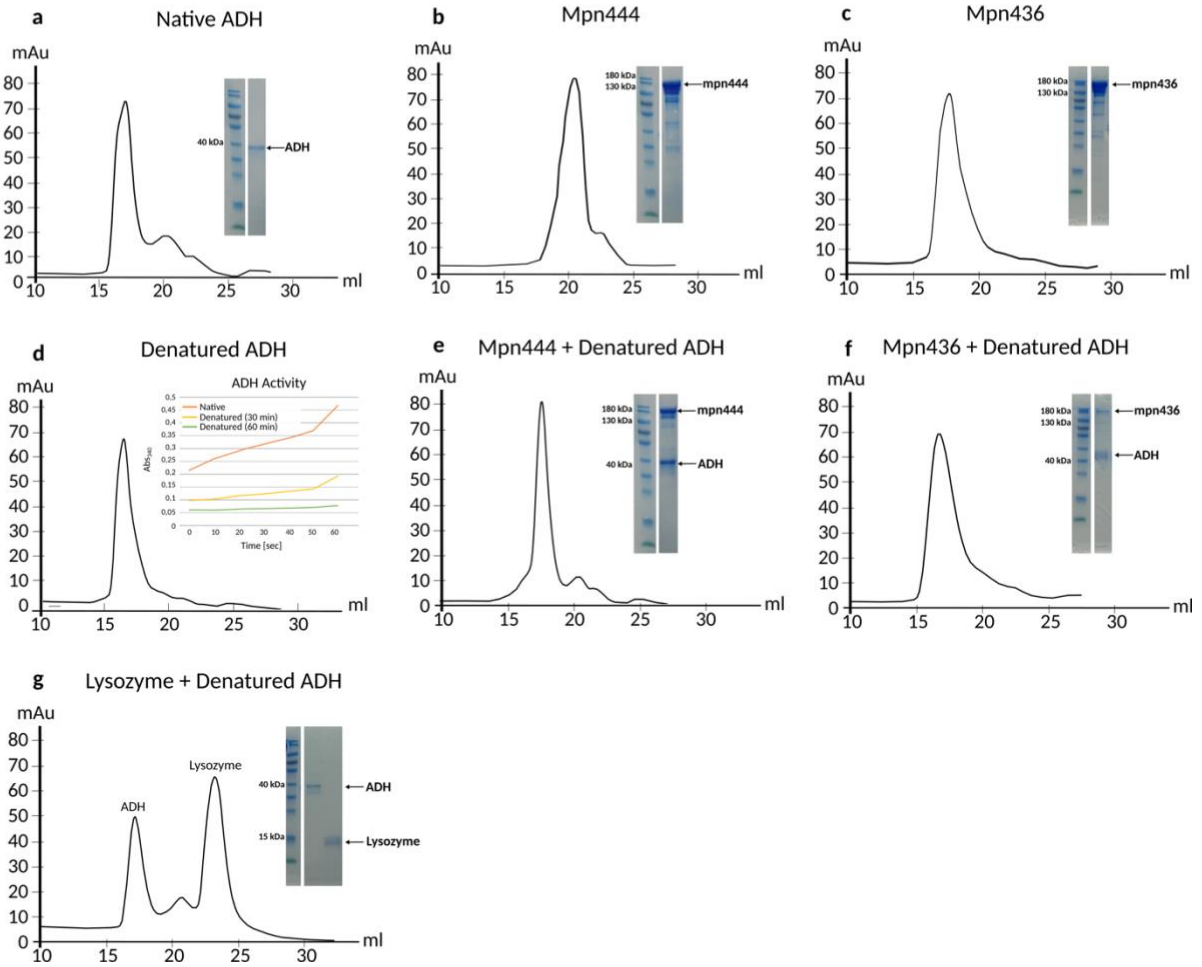

**Supplemental Figure 9: Chaperone-binding assay suggests that Mpn444 and Mpn436 interact with unfolded proteins**

The selective binding of Mpn444 and Mpn436 to unfolded client proteins was evaluated by size-exclusion chromatography (SEC). The proteins were eluted in Superose 6 Increase 10/300 GL column. The protein composition of the eluted fractions was assessed by SDS-PAGE.
**a** Control of the elution profile of native ADH
**b** Control of the elution profile Mpn444 with a molecular weight of approximately 140 kDa. Mpn444 was eluted around 21 ml.

**c** Control of the elution profile Mpn436 with a molecular weight of approximately 140 kDa. Mpn436 was eluted around 19 ml.
**d** Control of the elution profile of denatured ADH
**e** Chromatogram of Mpn444 and denatured ADH. The elution peak shifted from 21 to 17 ml. SDS gel confirms co-elution of Mpn444 and denatured ADH.
**f** Chromatogram of Mpn436 and denatured ADH. The elution peak shifted from 19 to 17 ml. SDS gel confirms co-elution of Mpn436 and denatured ADH
**g** Control for nonspecific binding. Lysozyme was used as a negative control. Lysozyme and denatured ADH did not co-elute.

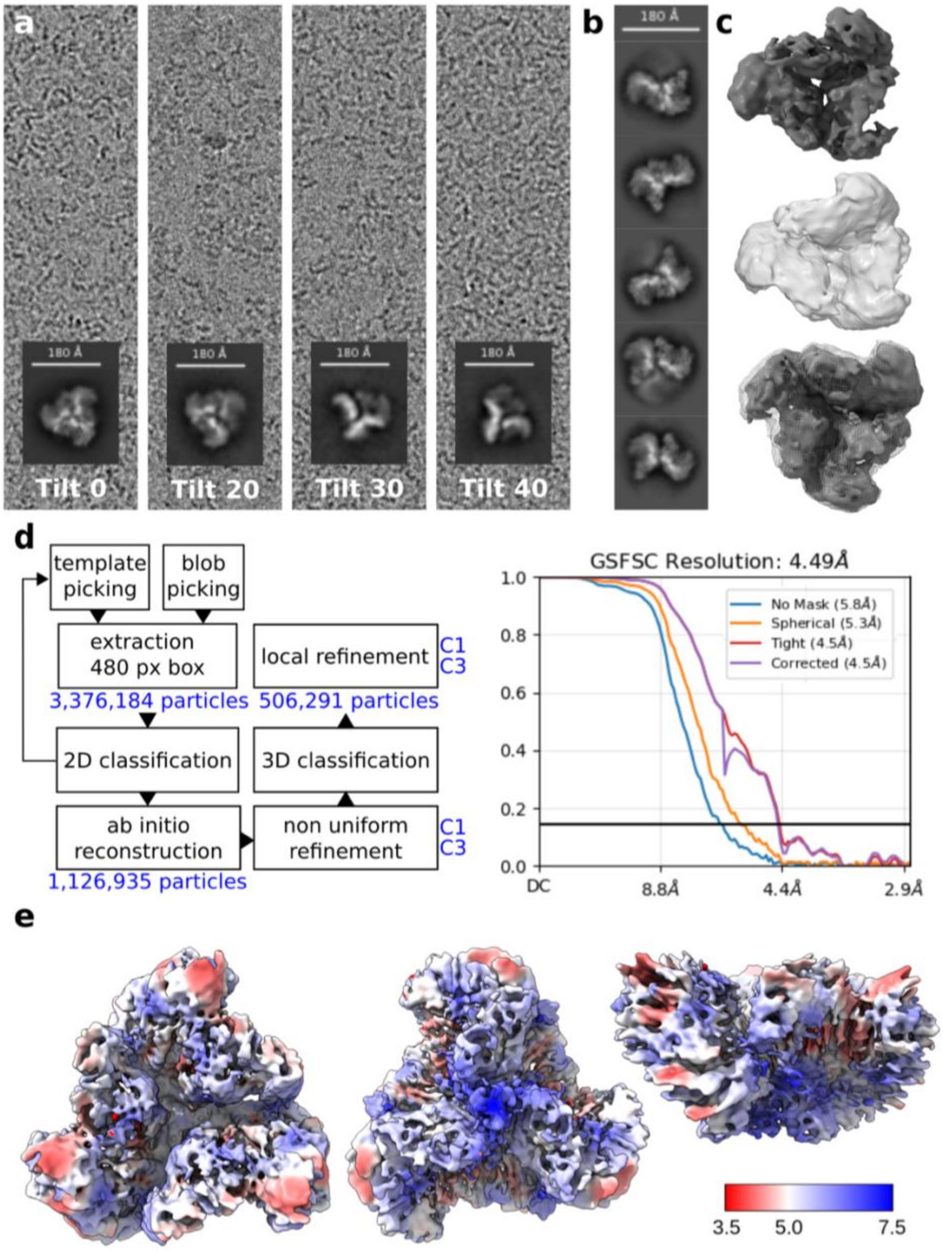

**Supplemental Figure 10: Processing pipeline of the Mpn444 trimer**
**a** Representative micrographs at 0.92 Å/pix with corresponding 2D classes (scale bar 180 Å).
**b** 2D classes of dimeric complexes (incomplete trimers) (scale bar 180 Å).

**c** *Ab initio* reconstruction of the Mpn444 trimer structure with C1 symmetry (dark grey, top) and C3 symmetry (light grey, middle). The superimposition of the *ab initio* reconstructions (bottom, C3 symmetry structure surface in mesh representation) shows that both have the same overall domain architecture. The density is weaker in one monomer in the C1 symmetry structure, which can be attributed to the persistent presence of monomers and dimers in the particle set after 2D classification.
**d** Processing pipeline of the Mpn444 trimer and FSC curve of the final reconstruction from cryoSPARC.
**e** Density map of the Mpn444 trimer at 5.4 Å, colored by local resolution, with the corresponding FSC curve from cryoSPARC.

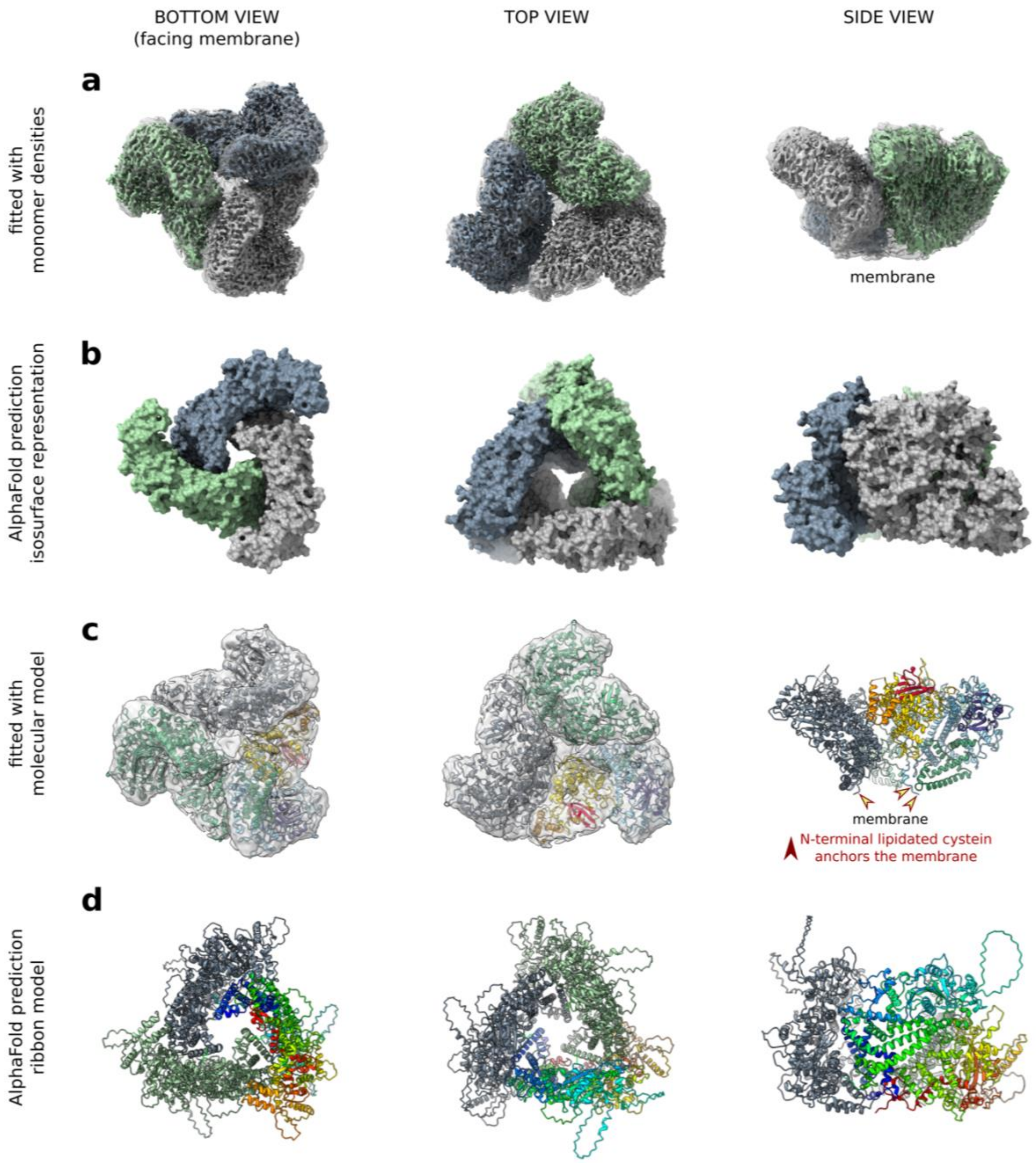

**Supplemental Figure 11: Comparison of the experimental trimer structure with the AlphaFold prediction**

**a** Density map of the Mpn444 trimer (transparent grey surface) fitted with the density maps of the Mpn444 monomer (dark blue, gray, sea green).

**b** Isosurface representation of the AlphaFold prediction of the Mpn444 trimer (dark blue, gray, sea green).

**c** Density map of the Mpn444 trimer (transparent grey surface) fitted with the molecular models of the Mpn444 monomer (ribbon models in dark slate grey, sea green, and a rainbow gradient from the N- to C-terminus). The N-terminal lipidated cysteines mediating membrane anchoring are indicated with red arrows.

**d** Ribbon model of the AlphaFold prediction of the Mpn444 trimer (dark slate grey, sea green, and a rainbow gradient from the N- to C-terminus).

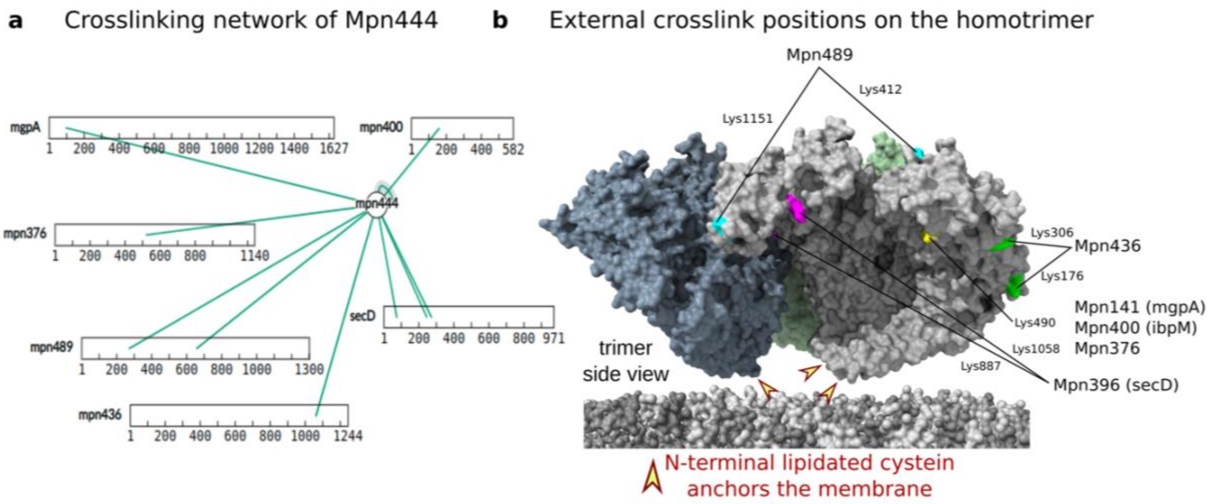

**Supplemental Figure 12: Positions of external crosslinks on the Mpn444 homotrimer**

**a** Mpn444 interacts with other *M. pneumoniae* proteins[14]: Mpn436, Mpn489, Mpn396 (secD), Mpn141 (mgpA/major adhesion complex), Mpn376, and Mpn400 (ibpM).

**b** Residues are colored according to the crosslinking partner: Lys412 and Lys1151 (cyan) interact with Mpn489; Lys176 and Lys306 (lime) interact with Mpn436; Lys887 and Lys1058 (magenta) interact with Mpn396 (SecD); and Lys490 (yellow) interacts with Mpn141 (mgpA/major adhesion complex), Mpn376, and Mpn400 (ibpM).

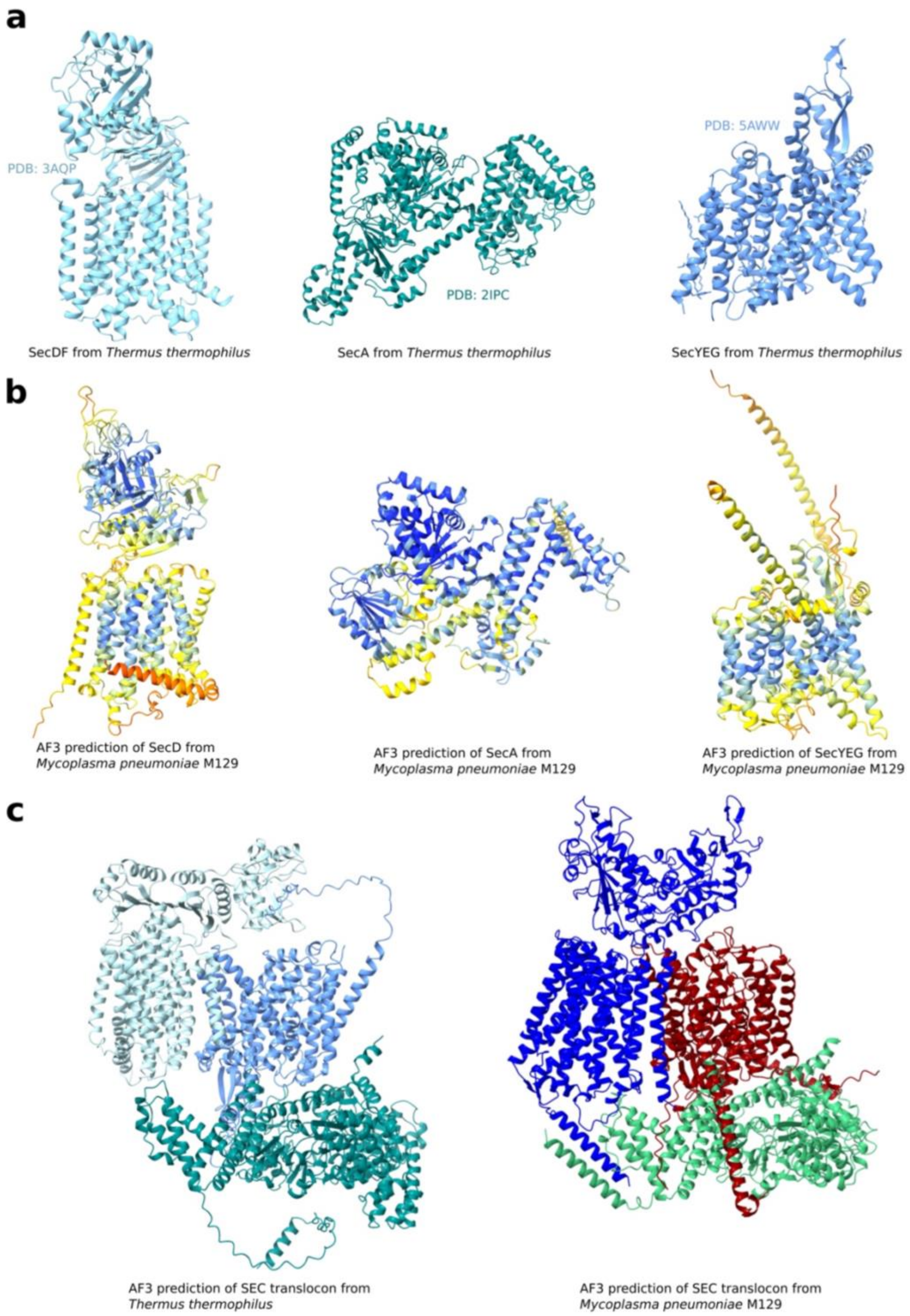

**Supplemental Figure 13: Structural comparison of the SEC translocon from *M. pneumoniae* and *Thermus thermophilus*.**

**a** Experimental structures of the proteins SecDF (left, PDB: 3AQP, light blue), SecA (middle, PDB: 2IPC, sea green) and SecYEG (right, PDB: 5AWW, blue) forming the SEC translocon in *T. thermophilus*.

**b** AlphaFold3 predictions of the proteins SecD, SecA and SecYEG forming the SEC translocon in *M. pneumoniae.* The color scheme represents the confidence level of the prediction from AlphaFold (dark blue for very high confidence, light blue for high confidence, yellow for low confidence, dark orange for very low confidence).

**c** AlphaFold3 predictions of the SEC translocon from *T. thermophilus* (left) and *M. pneumoniae.* The colors are as in (a) for *T. thermophilus* and as in Figure 4 for *M. pneumoniae.*

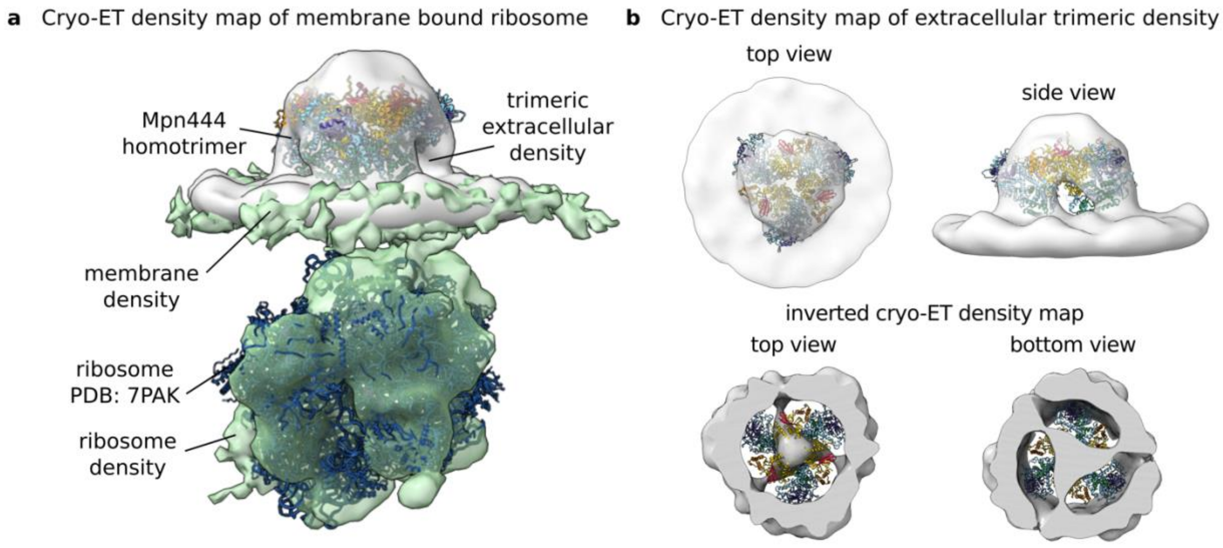

**Supplemental Figure 14: Sub-tomogram average showing co-localization of a trimeric density extracellularly associated with the ribosome**

**a** Superimposed cryo-ET density maps of membrane-bound ribosome (in green) and extracellular trimeric density (in grey). Both maps were created with the same particle set (Cryo-ET Data Portal, dataset ID: DS-10003), but one was aligned on the ribosome (green, mask including ribosome and membrane) and the other on the extracellular density (grey, mask including extracellular density and membrane). The green map was fitted with a ribbon model of the Mycoplasma ribosome (PDB: 7PAK) and the grey map was fitted with the ribbon model of the homotrimeric Mpn444 assembly.

**b** Cryo-ET density maps of extracellular trimeric density. Shown in top and side view (top) as well as inverted in top and bottom view (bottom). The density map is fitted with the ribbon model of the homotrimeric Mpn444 assembly.
